# Metabolic constraints on nitrogen fixation by rhizobia in legume nodules

**DOI:** 10.1101/2021.02.16.431433

**Authors:** Carolin C. M. Schulte, Khushboo Borah, Rachel M. Wheatley, Jason J. Terpolilli, Gerhard Saalbach, Nick Crang, Daan H. de Groot, R. George Ratcliffe, Nicholas J. Kruger, Antonis Papachristodoulou, Philip S. Poole

## Abstract

Rhizobia induce nodule formation on legume roots and differentiate into bacteroids, which use plant-derived dicarboxylates as energy and electron sources for reduction of atmospheric N_2_ into ammonia for secretion to plants. Using heterogeneous genome-scale datasets, we reconstructed a model of bacteroid metabolism to investigate the effects of varying dicarboxylate and oxygen supply on carbon and nitrogen allocation. Modelling and ^13^C metabolic flux analysis in bacteroids indicate that microaerobiosis restricts the decarboxylating arm of the TCA cycle and limits ammonia assimilation into glutamate. Catabolism of dicarboxylates induces a higher oxygen demand but also a higher NADH/NAD^+^ ratio compared to sugars. Carbon polymer synthesis and alanine secretion by bacteroids facilitate redox balance in microaerobic nodules with alanine secretion increasing as oxygen tension decreases. Our results provide a framework for understanding fundamental constraints on rhizobial metabolism during symbiotic nitrogen fixation.

## Introduction

Biological nitrogen fixation provides 50 – 70 Tg of bioavailable nitrogen in agricultural systems per year^1^ and sustains global food security. The most efficient contribution to biologically fixed nitrogen is from symbioses between legumes and rhizobia^2^, which are soil bacteria that induce formation of nodules on plant roots. Inside nodules, rhizobia differentiate into bacteroids that reduce atmospheric N_2_ into ammonia for secretion to the plant host in exchange for dicarboxylates, primarily succinate and malate^3,4^. Succinate and malate are typically metabolised via malic enzyme and pyruvate dehydrogenase, yielding acetyl-CoA that can be oxidised in the tricarboxylic acid (TCA) cycle^5^. Whether bacteroids need a complete TCA cycle remains unclear, as it is essential for *Rhizobium* and *Sinorhizobium* species, but mutants of *Bradyrhizobium japonicum* lacking 2-oxoglutarate dehydrogenase activity achieve wild-type levels of N_2_ fixation on a per bacteroid basis^6,7^. Flux through the TCA cycle produces NADH and FADH_2_, which provide electrons for nitrogen fixation and ATP generation^8^. Despite rhizobia requiring oxygen for ATP synthesis, oxygen levels in nodules are only 10 – 40 nM^9^, which is a requirement for highly oxygen-sensitive nitrogenase enzyme activity^10^. The challenge of balancing carbon allocation under these conditions is evidenced by the synthesis of lipids and carbon polymers, such as polyhydroxybutyrate (PHB), which indicate imbalances in nutrient supply^8,11^. PHB and lipids have been suggested to play a role as carbon and redox sinks for bacteroids, although their accumulation is variable between rhizobial strains and their role remains to be elucidated^8,12,13^.

The defining distinction between N_2_ fixation by rhizobial bacteroids compared to free-living bacteria is the secretion of fixed ammonia to the plant. However, there is no known metabolic mechanism forcing secretion of fixed nitrogen to the plant instead of assimilation by the bacteroid. In addition to the main secretion product ammonia, a significant portion of fixed nitrogen is apparently secreted in the form of alanine and aspartate^14,15^. At the same time, ammonia assimilation by the glutamine synthetase-glutamine oxoglutarate aminotransferase (GS-GOGAT) system, which is active in free-living rhizobia, is downregulated during the symbiosis^16–18^.

Due to the complexity of bacteroid metabolism, several studies have used computational approaches to investigate the symbiosis. Notably, metabolic models for various rhizobial species have been reconstructed^19–22^, and some reconstructions have been integrated with models for the host plant^23,24^. Metabolic models describe all enzymatic and transport reactions encoded in an organism’s genome and enable the simulation of metabolic flux distributions under defined environmental conditions^25,26^. The most common approach for analysing metabolic models is flux balance analysis (FBA), where an objective function reflecting the evolutionary goal of the organism under investigation is optimised to calculate a flux distribution^27^. Maximum cellular growth is the most commonly used objective function and has been found to reproduce experimental results for strains adapted to growth under laboratory conditions^28^, but this is clearly not applicable in the case of growth-arrested bacteroids. *In silico* studies of bacteroid metabolism, which have so far mostly applied standard FBA methods, defined an objective reaction comprising nitrogen export to the plant and synthesis of storage compounds^19–22^.

The result of FBA calculations is a single flux distribution, which is often not a unique solution for the optimisation problem^27^. In contrast, methods such as elementary flux mode enumeration describe all minimal sets of steady state fluxes through the metabolic network^29^. While this approach is significantly more computationally expensive than FBA and currently not feasible for genome-scale metabolic networks^30^, it provides a more comprehensive and unbiased description of metabolism that does not rely on an artificially defined objective. Addressing the problem of combinatorial explosion during elementary flux mode enumeration, elementary conversion modes (ECMs) have been proposed as an alternative approach to compute all possible stoichiometries between input and output metabolites^31^. Although ECMs do not provide information on the underlying metabolic pathways for a specific conversion, they can still capture all metabolic capabilities of an organism. The feasibility of ECM enumeration for genome-scale metabolic networks when focusing on subsets of metabolites has recently been demonstrated^32^.

Despite decades of research efforts, a comprehensive view of the links between central carbon and nitrogen metabolism in bacteroids is missing. Most experimental studies have focused on individual metabolic pathways, while computational models have employed artificial objective functions that may not be relevant in the natural system.

In this study, we combine experimental and metabolic modelling approaches to explain fundamental features of bacteroid metabolism. Using proteome, transcriptome and gene essentiality data, we reconstruct a model of metabolic pathways active during nitrogen fixation in *Rhizobium leguminosarum* bv. *viciae*. We implement modelling strategies that circumvent the limitations of traditional FBA to explain the importance of the experimentally observed storage polymer synthesis and amino acid secretion in bacteroids. We further investigate the role of the TCA cycle during nitrogen fixation and validate model predictions by ^13^C metabolic flux analysis of *Azorhizobium caulinodans*. Our model provides insights into the fundamental constraints on rhizobial metabolism during symbiotic nitrogen fixation. An improved understanding of metabolic processes in bacteroids is of central importance for ongoing efforts in optimising existing symbioses and engineering novel plant-microbe interactions for sustainable agriculture^2,33^.

## Results

### Data-based reconstruction of a bacteroid metabolic model

We reconstructed a metabolic model for pea bacteroids of *Rhizobium leguminosarum* bv. *viciae* 3841 using bacteroid-specific experimental data. First, we quantified the proteome of unlabelled bacteroids relative to ^15^N-labelled free-living bacteria (Fig. 1, Supplementary Fig. 1, Supplementary Note 1, Supplementary Data 1). In addition, genes upregulated in transcriptional datasets of bacteroids^34^ (Supplementary Fig. 2) and genes identified as specifically essential for symbiosis formation by insertion sequencing (INSeq)^35^ were included in the model. The final core metabolic network named *i*CS323 contained 323 genes, 237 metabolites and 299 reactions, 207 of which are metabolic (excluding transport, demand and sink reactions) (Supplementary Table 1, Supplementary Data 2-4). Out of the 207 metabolic reactions, 177 (86%) are supported by experimental evidence from at least one of the bacteroid-specific datasets. *i*CS323 was evaluated using MEMOTE^36^, confirming the absence of stoichiometrically balanced cycles, orphan metabolites and dead-end reactions. (Supplementary Data 5).

**Figure 1.**
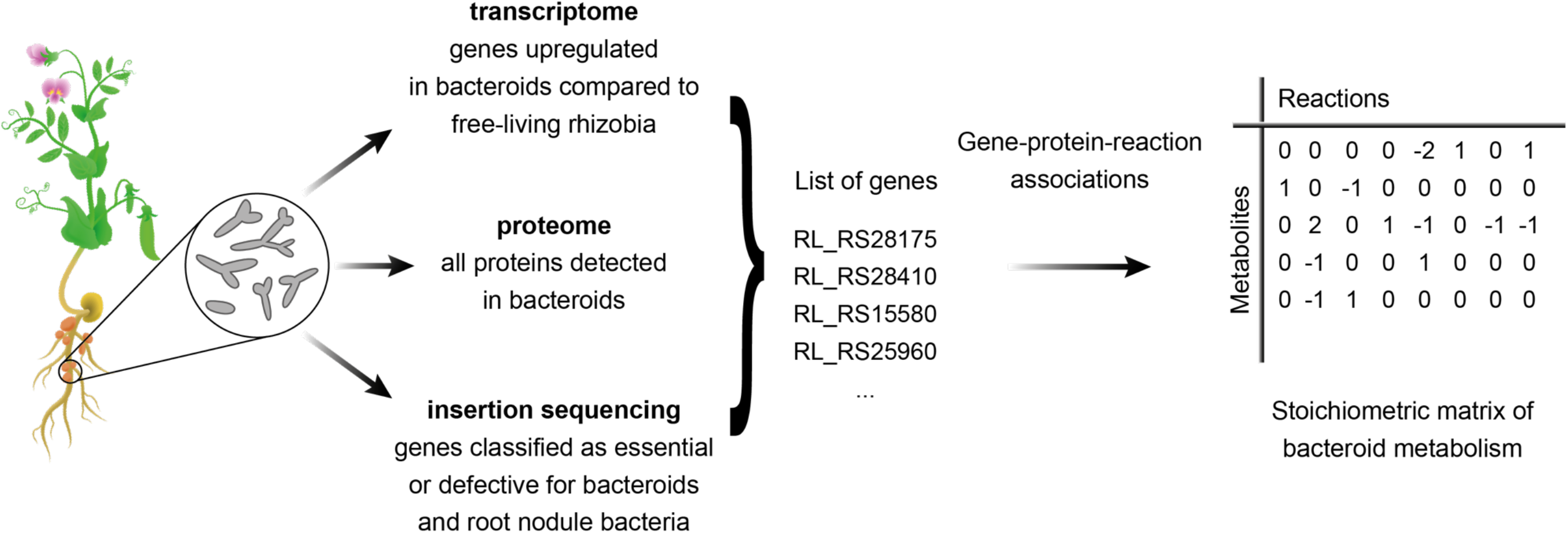
Workflow for metabolic model reconstruction. A metabolic model of *R. leguminosarum* bacteroids was reconstructed using transcriptome, proteome and gene essentiality data.

The main pathways in the final model were central carbon metabolism (TCA cycle, gluconeogenesis, pentose phosphate pathway), amino acid metabolism and carbon polymer synthesis. As an initial validation, malate, succinate and oxygen uptake were constrained according to the flux boundaries defined in a modelling study of *Sinorhizobium meliloti*^21^, and standard FBA maximising nitrogenase activity was performed. The obtained flux distribution captured key features of bacteroid metabolism, including use of the TCA cycle, pyruvate synthesis via malic enzyme and minor activity of gluconeogenesis^5,8,37^ (Supplementary Fig. 3).

When comparing gene essentiality predictions to mutant phenotypes identified by INSeq^35^, we found that 280 (87%) of the genes in the model had an *in silico* phenotype that agreed with the experimental evidence (Supplementary Data 6). Genes were defined as essential *in silico* if their deletion prevented flux through the nitrogenase reaction in the model. Of the 31 false negatives (*in silico* non-essential genes that were essential according to INSeq), nine were involved in sugar metabolism. In agreement with the results of a recent modelling study of *S. meliloti*^24^, this may suggest that rhizobia differentiating into bacteroids have access to sugars as a carbon source, whereas nitrogen-fixing bacteroids do not. In addition, four genes involved in serine metabolism were incorrectly predicted to be non-essential. A possible explanation is the role of serine as a precursor for cysteine biosynthesis. Cysteine plays a role in the synthesis of iron-sulphur clusters for the nitrogenase enzyme^38^, which is not explicitly accounted for in the model. Pyruvate kinase was further predicted to be non-essential in disagreement with the INSeq data. A *pykA* mutant had higher nitrogenase activity than the wild-type in plants harvested after 28 days^5^, indicating that the INSeq phenotype may be a result of developmental delay and the model correctly identifies the gene as non-essential.

Overall, our model shows good predictive quality for gene essentiality. It is important to note that perfect agreement of *in silico* predictions with the experimental data is not expected. This is due to our model being limited to central metabolic pathways in bacteroids, neglecting for example the synthesis of nucleotides for DNA replication. In addition, genes that are determined to be essential for bacteroids by INSeq may actually cause a growth defect at earlier stages of symbiosis formation and are not necessarily required for nitrogen fixation itself.

### Characterisation of bacteroid metabolism using elementary conversion modes

Previously published modelling studies of bacteroid metabolism used FBA and maximised a lumped objective reaction comprising ammonia and amino acid export as well as storage polymer synthesis^19–22^. While this approach constrains flux distributions to reflect experimentally observed phenotypes, it also introduces an artificial stoichiometric coupling between the objective metabolites, which precludes the investigation of changes in carbon and nitrogen allocation depending on nutrient availability. To characterise bacteroid metabolism with a minimum number of pre-set assumptions, we used ecmtool^32^ to enumerate ECMs of the metabolic network (Fig. 2, Supplementary Data 7). ECMs capture all metabolic capabilities in terms of input-output stoichiometry and therefore provide a comprehensive description of metabolism without assuming optimality with respect to a specific objective.

**Figure 2.**
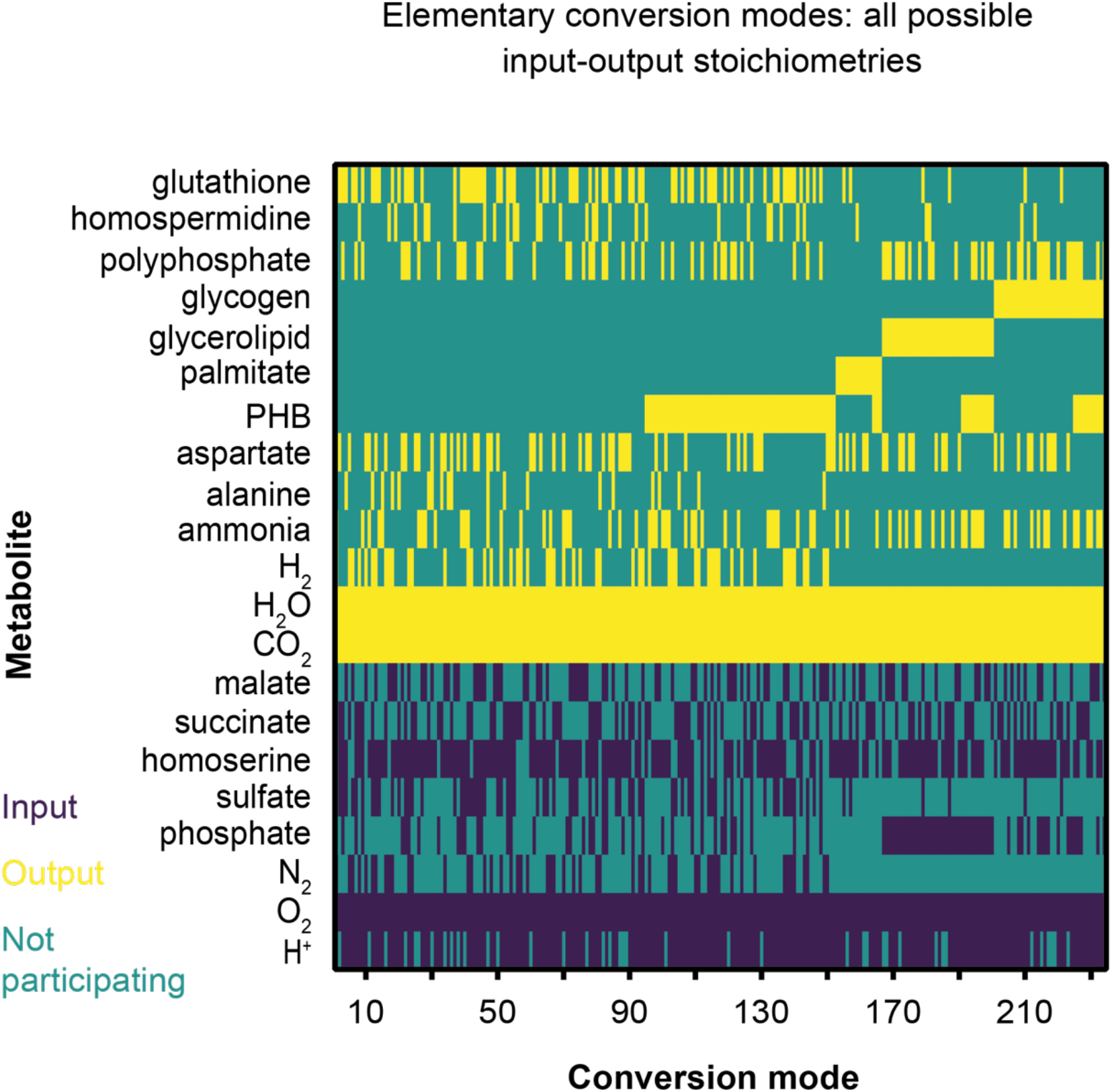
Elementary conversion modes for *i*CS323. The heatmap represents conversion modes calculated with succinate and malate as carbon sources and carbon polymers (palmitate, PHB, glycerolipid, glycogen), ammonia, alanine and aspartate as outputs. Flux values were indexed to represent inputs, outputs and non-participating compounds.

Conversion modes were found for all expected products (ammonia, alanine, aspartate and storage compounds) with a minimal set of inputs, mainly malate/succinate, oxygen and N_2_ (Supplementary Data 8). The presence of conversion modes for storage polymer synthesis without nitrogenase activity suggests that the two processes are not inherently linked, which agrees with reports of storage polymer accumulation prior to the onset of nitrogen fixation^12,39^. Potential co-catabolism of an amino acid in addition to malate/succinate was investigated by analysing conversion modes with each of the proteinogenic amino acids or *γ*-aminobutyric acid (GABA) as an additional input. Conversion modes were found for arginine, cysteine, GABA, glutamine, glutamate, glycine, serine and threonine. All conversions involving arginine, cysteine or serine had a negative net nitrogen output and were therefore considered unlikely to be biologically relevant. For the remaining conversion modes with a positive net nitrogen output, no benefit was found for any amino acid in terms of oxygen demand or carbon cost (Supplementary Fig. 4). We therefore limited our analysis to conversion modes using GABA, which is known to be metabolised in pea bacteroids^40^.

Oxygen demand per carbon uptake was decreased for all conversion modes that produced storage polymers compared to conversions that did not (Fig. 3a, Supplementary Fig. 5). Carbon storage polymers thus function as carbon and redox sinks under oxygen-limiting conditions, enabling electrons to be partitioned to polymer synthesis rather than oxygen as a terminal electron acceptor. The oxygen uptake relative to fixed N_2_ was increased for conversion modes generating glycogen or lipids (Fig. 3b), consistent with the ATP requirement for producing these storage compounds, which would add to the ATP demand of the nitrogenase reaction. In addition, conversion modes generating carbon polymers increased the carbon cost per nitrogen supplied to the plant (Fig. 3c), indicating that storage compounds divert resources from nitrogen fixation. This aligns with observations that plant nodule cells accumulate starch when occupied by bacteroid glycogen synthase mutants^12^, i.e. the plant has excess carbon when bacteroid polymer synthesis is restricted.

**Figure 3.**
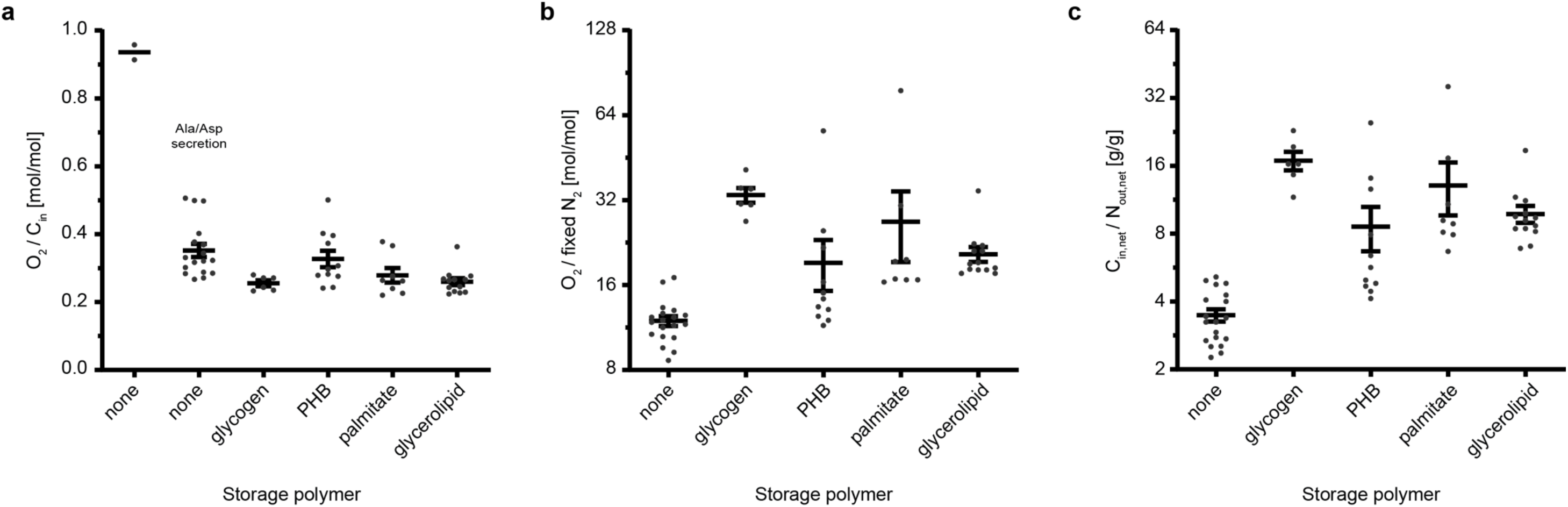
Effect of polymer synthesis and amino acid secretion on oxygen demand and carbon cost of nitrogen fixation. **a** Oxygen uptake per carbon uptake, **b** oxygen uptake per fixed N_2_ and **c** carbon cost (difference of carbon input and output) per nitrogen secreted (difference of nitrogen input and output) were determined for conversion modes using succinate, malate and GABA as carbon sources. In **a**, ECMs without storage polymer production have been separated into those secreting only ammonia and those secreting alanine and/or aspartate in addition to/instead of ammonia. Each data point represents an individual conversion mode, lines and bars indicate mean ± SEM.

The carbon cost determined for conversion modes without polymer production is similar to the theoretical cost of nitrogen fixation (2.5 g carbon/g nitrogen), whereas that for PHB- and some palmitate- and glycerolipid-producing conversion modes is close to the experimentally observed value (8 g carbon/g nitrogen)^2^. Given that the plant host regulates the supply of nutrients such as carbon and oxygen in response to nitrogen output^41,42^, this would potentially limit excess diversion of carbon into polymers.

Conversions without carbon polymer production could be sub-grouped into those generating alanine and/or aspartate and those only producing ammonia, where amino acid secretion reduced oxygen demand per carbon uptake (Fig. 3a) and oxygen demand per nitrogen output (Supplementary Fig. 5). Alanine dehydrogenase catalyses the NADH-dependent synthesis of alanine from pyruvate and ammonia, making this pathway an oxygen-independent carbon sink for NAD^+^ regeneration as well as a sink for carbon and protons. Synthesis of alanine in particular enables bacteroids to maintain nitrogen export to the plant in a low oxygen environment and explains the mixed secretion of ammonia and amino acids observed experimentally^14^. Amino acid secretion could be favoured over polymer synthesis under certain conditions since it allows for removal of carbon from the bacteroid rather than intracellular accumulation.

The same trends for storage polymer synthesis and amino acid secretion were observed for ECMs of a model for *Sinorhizobium fredii* bacteroids^19^ (Supplementary Fig. 6), indicating that these principles are likely to govern symbiotic metabolism across rhizobial species.

### Role of oxygen limitation in shaping bacteroid metabolism

We next characterised the network response to different carbon and oxygen availability when optimum nitrogenase activity is maintained. Since uptake fluxes of bacteroids are difficult to determine experimentally, we performed phenotype phase plane analysis^43^. Avoiding artificial biases of carbon and nitrogen allocation, nitrogenase activity was evaluated without maximising the production of storage compounds. Four feasible regions with distinct metabolic behaviour were identified, with phase I characterised by carbon limitation and increasing oxygen limitation from phase II to phase IV (Fig. 4a, Supplementary Fig. 7).

**Figure 4.**
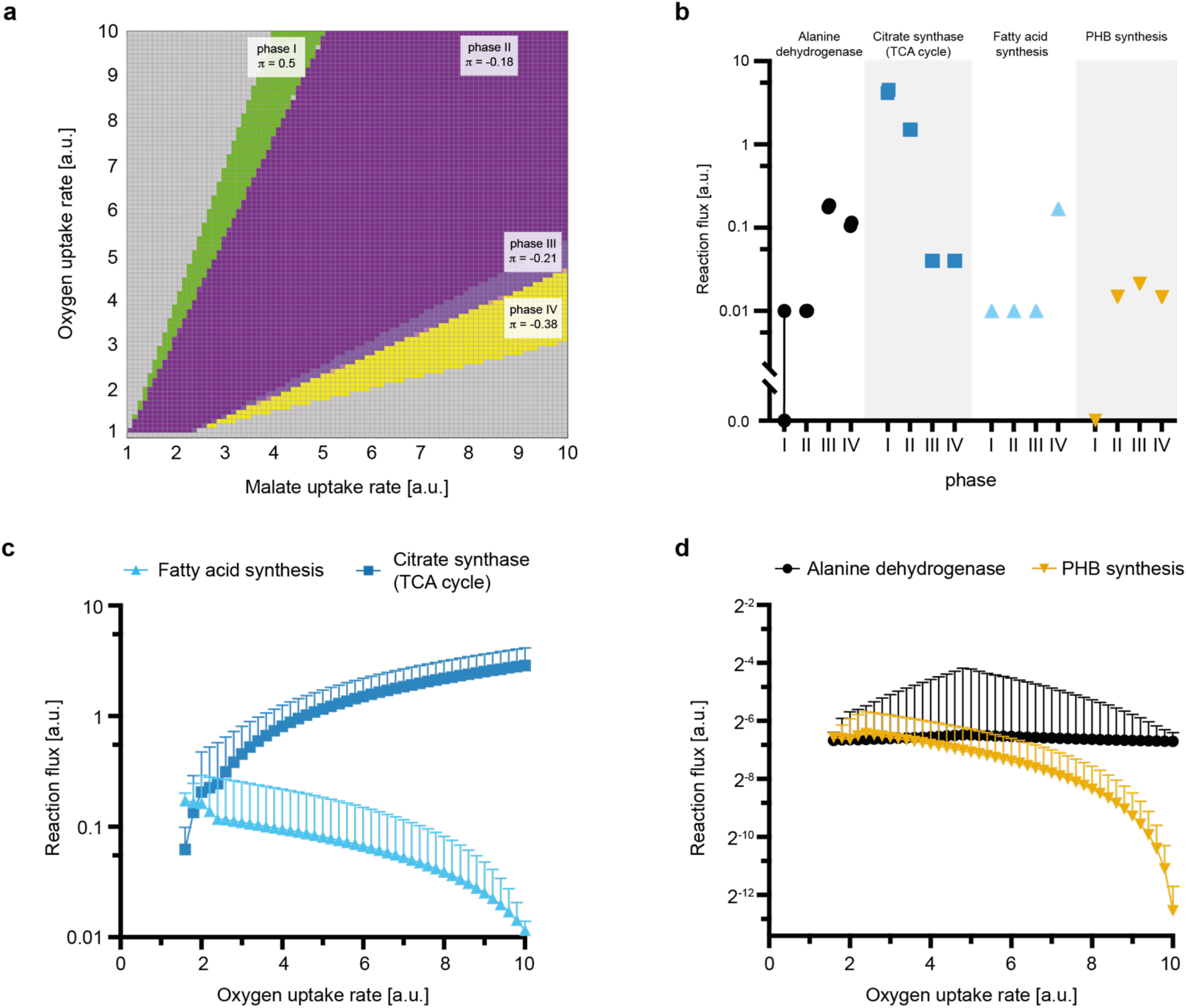
Metabolic response of bacteroids to varying carbon and oxygen availability. Phenotype phase plane analysis with varying malate and oxygen uptake rates was performed for *i*CS323. **a** Shadow prices (*π*) for oxygen in the four phases defined by phenotype phase plane analysis. **b** Flux ranges for alanine dehydrogenase, citrate synthase, fatty acid and PHB synthesis fluxes determined by flux variability analysis for the four phases defined in **a**. Symbols indicate upper and lower bounds of the indicated flux for maximum nitrogenase activity. Note the symbols for upper and lower bounds are overlapping in most cases. **c** Fatty acid synthesis and citrate synthase activity at a fixed malate uptake rate of 4 flux units and varying oxygen uptake rates were determined by ensemble-evolutionary FBA. **d** Same as **c** for PHB synthesis and alanine dehydrogenase. Values plotted in **c** and **d** represent mean values and standard deviation for 50,000 random objective functions.

Flux variability analysis showed that with increasing oxygen limitation, flux through TCA cycle enzymes, as represented by citrate synthase, decreased (Fig. 4b). Carbon was increasingly channelled into pyruvate, which caused accumulation of PHB and lipids as well as alanine production, with all nitrogen being secreted in the form of alanine instead of ammonia in phase IV. The tendency to increase alanine secretion at low oxygen levels has recently been shown experimentally in *B. japonicum*^44^. Alanine secretion is thus important to sustain bacteroid metabolism under oxygen limitation, which probably occurs in the natural system^45^. This is consistent with a 20% reduction in dry weight of peas inoculated with alanine dehydrogenase mutants of *R. leguminosarum* that no longer secrete alanine^46^.

Independently of maximum nitrogenase activity, general network properties were investigated using ensemble-evolutionary FBA^47^. The results supported increased PHB and lipid synthesis under oxygen-limiting conditions as well as decreased activity of the TCA cycle (Fig. 4c, d). The trend for increased alanine synthesis was less clear, indicating that it is linked to nitrogen fixation rather than being an inherent network property. The observed shifts in metabolic behaviour support the role of alanine, PHB and lipids as sinks for carbon when oxygen is limiting, and usage of the TCA cycle becomes disadvantageous due to accumulation of reduced electron carriers^8^, which agrees with the conversion mode analysis.

### Metabolic constraints on ammonia assimilation by bacteroids

The predicted downregulation of the TCA cycle would reduce the availability of 2-oxoglutarate required by the GS-GOGAT pathway, which is the sole pathway for ammonia assimilation coupled to growth in rhizobia. It should be noted that rhizobial GS-GOGAT mutants cannot grow on ammonia as a nitrogen source, and alanine dehydrogenase for example cannot substitute for GS-GOGAT^18^. Increased ammonia assimilation into glutamate by bacteroids would require increased TCA cycle activity and hence increase oxygen demand as indicated by a significant positive correlation between oxygen uptake and glutamine synthetase activity in the conversion mode analysis (Fig. 5a). To assess the impact of ammonia assimilation on the metabolic fluxes in bacteroids, we forced flux through a demand reaction for glutamate in the model. In this scenario, flux through the decarboxylating arm of the TCA cycle had to be maintained under oxygen limitation to supply 2-oxoglutarate for ammonia assimilation, which caused an increased oxygen demand (Fig. 5b).

**Figure 5.**
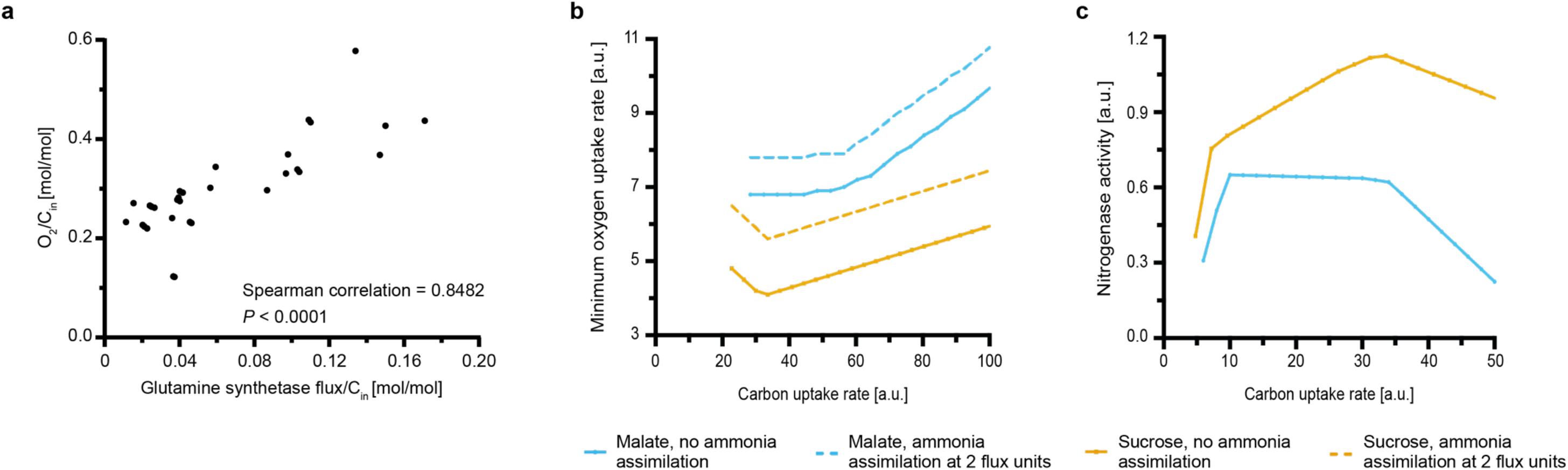
Predicted impact of ammonia assimilation on oxygen demand. **a** Scatterplot showing the relationship between oxygen demand and glutamine synthetase activity per carbon uptake predicted by ECM analysis. Each point represents an individual conversion mode, only conversion modes with glutamine synthetase activity are shown. **b** Predicted minimum oxygen demand for malate or sucrose as a carbon source without (solid line) and with (dashed line) ammonia assimilation by bacteroids. Oxygen demand is shown for equivalent nitrogenase activity for the respective carbon source. **c** Predicted maximum nitrogenase activity for malate or sucrose as a carbon source and a maximum oxygen uptake rate of 4 flux units. Values in **b** and **c** are shown per mol of carbon atoms.

Our modelling results indicate the importance of the carbon source provided to bacteroids to enforce use of the TCA cycle as the main catabolic pathway. It is currently unclear why plants provide bacteroids with C4-dicarboxylates, especially considering that photosynthate is transported to nodules as sucrose^48^. We therefore compared the effects of utilising sucrose instead of malate as a carbon source *in silico*. For a given carbon uptake rate, the model predicted higher nitrogenase activity for sucrose compared to malate (Fig. 5c) in agreement with an integrated plant-bacteroid model for *S. meliloti*^24^. Furthermore, less oxygen was needed for sucrose (Fig. 5b) (or glucose, Supplementary Fig. 8) catabolism, which is in accordance with experimentally determined oxygen uptake rates of free-living *R. leguminosarum* (Fig. 6a). Catabolism of arabinose, a sugar metabolised via 2-oxoglutarate, induced an oxygen demand similar to growth on succinate. This suggests that catabolism of TCA cycle intermediates generally creates a high oxygen demand and thus causes a more severe growth impairment compared to glucose under low oxygen conditions (Fig. 6c). However, the NADH/NAD^+^ ratio was similar for growth on glucose and arabinose, but significantly higher for succinate catabolism (Fig. 6b). This could be explained by 2-oxoglutarate supply from arabinose catabolism partly obviating use of the decarboxylating TCA cycle arm, but also by differences in the regulation of sugar versus dicarboxylate uptake and metabolism.

**Figure 6.**
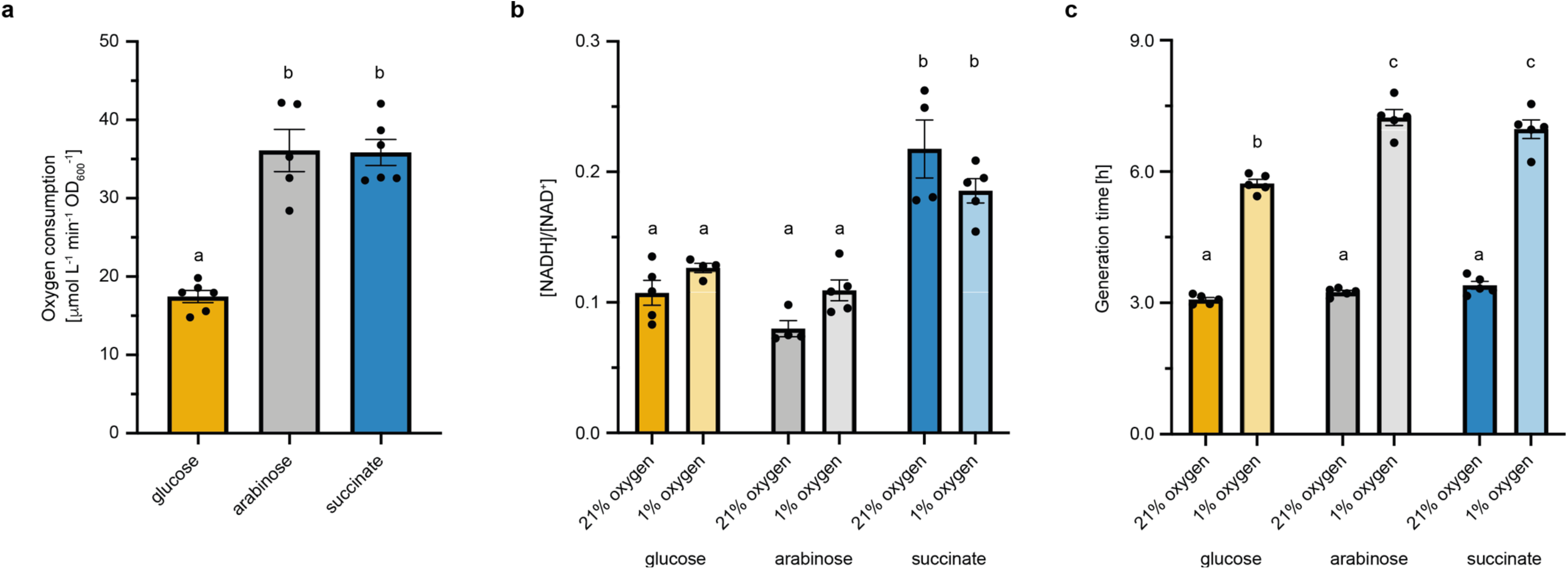
Carbon source determines oxygen demand and redox ratio in *R. leguminosarum*. **a** Experimentally determined oxygen consumption rate of *R. leguminosarum* grown in minimal media with NH_4_Cl as a nitrogen source and glucose, arabinose or succinate as the sole carbon source. **b** NADH/NAD^+^ ratio and **c** generation time were measured for cultures grown at 21% oxygen and 1% oxygen. Data points represent independent biological replicates with lines and error bars indicating mean ± SEM. Lowercase letters indicate significant differences between samples determined by one-way ANOVA followed by Tukey’s multiple comparisons test.

Overall, these findings indicate that supply of dicarboxylates such as succinate to bacteroids is particularly suitable for creating both a high oxygen demand and a highly reduced redox ratio, which is important for supply of electrons to the nitrogenase enzyme^8^. The absolute value of the redox ratio did not change significantly for different oxygen concentrations and was also not correlated with the growth rate (Fig. 6c). Thus, the redox state of *R. leguminosarum* is mainly determined by the carbon source, implying that the nature and quantity of the supplied carbon source are of central importance for defining metabolic fluxes in bacteroids, with succinate providing a higher NADH/NAD^+^ ratio than glucose. Remarkably, plants provide dicarboxylates as a carbon source to bacteroids even though they are less efficient at supporting N_2_ fixation and increase oxygen demand relative to sucrose in the oxygen-limited nodule.

### Metabolic flux analysis of *Azorhizobium caulinodans* supports model predictions

To validate core findings of our model predictions experimentally, we performed ^13^C metabolic flux analysis of *Azorhizobium caulinodans* ORS571. We chose this rhizobial strain for its ability to perform nitrogen fixation in both free-living conditions and in symbiosis with *Sesbania rostrata*. When comparing non-diazotrophic growth of *A. caulinodans* at different oxygen levels, PHB synthesis increased 3-fold under microaerobic (3% O_2_) conditions compared to aerobic growth, and further increased in bacteroids (Fig. 7, Supplementary Data 9). This agrees with the predicted importance of storage polymer synthesis for balancing carbon allocation under oxygen-limited conditions.

**Figure 7.**
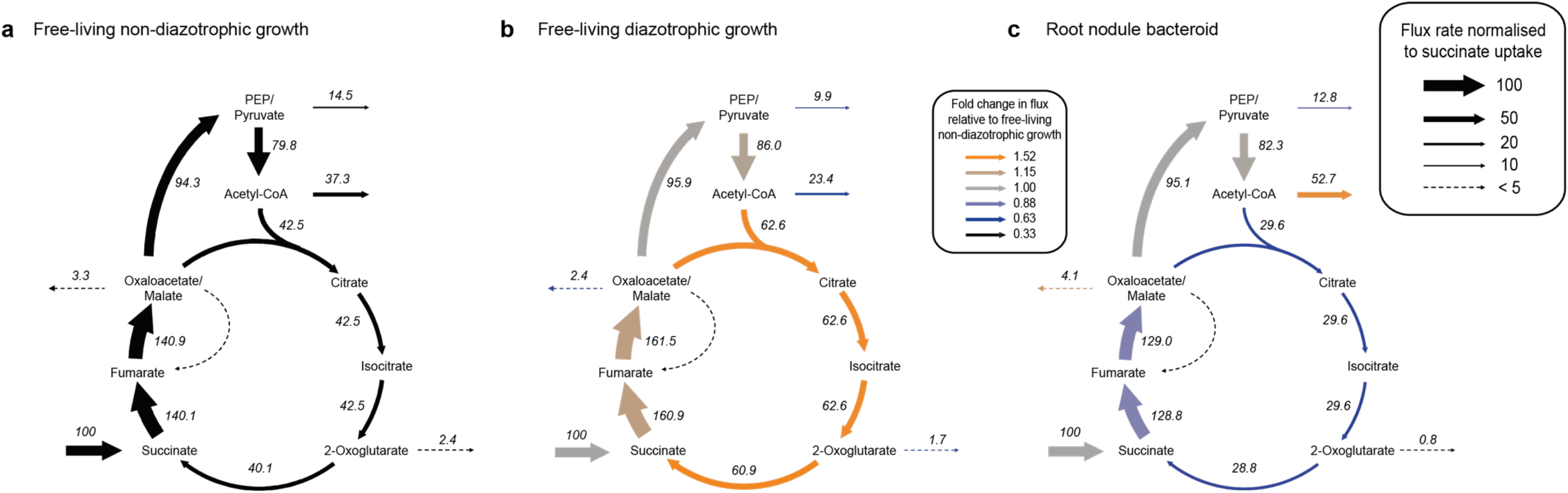
Metabolic flux analysis of *Azorhizobium caulinodans*. Labelling experiments were performed with [^13^C_4_]-succinate for *A. caulinodans* **a** grown in free-living conditions with NH_4_Cl as a nitrogen source, **b** grown in free-living conditions without a nitrogen source and **c** bacteroids isolated from root nodules of *Sesbania rostrata*. Flux values were normalised to a succinate uptake rate of 100 units. Colours in **b** and **c** indicate fold changes in flux values compared to **a**. Note the flux from succinate to fumarate is the sum of the succinate uptake rate and the flux through the decarboxylating arm of the TCA cycle.

TCA cycle fluxes increased during free-living diazotrophic growth compared to non-diazotrophic growth, which is consistent with an increased demand for reductant for nitrogen fixation as well as for cell growth. However, an overall downregulation of the TCA cycle was observed when comparing bacteroid metabolism to non-diazotrophic growth under microaerobic conditions. Importantly, the highest relative downregulation was observed for the decarboxylating arm of the TCA cycle between citrate and succinate. As predicted by our model, bacteroids would thus have limited capability for 2-oxoglutarate synthesis and would consequently be restricted for ammonia assimilation into glutamate.

## Discussion

In this study, we combined experimental and computational methods to provide explanations for fundamental properties of bacteroid metabolism, which have so far mostly been investigated separately.

Our modelling results indicate that oxygen limitation is the driving factor behind the observed synthesis of carbon polymers^8,12^ and induces the secretion of alanine, in addition to ammonia. The level of alanine secretion is dependent on the ratio of carbon and oxygen supply, which agrees with experimental studies^44^. Rhizobial nitrogen fixation is fuelled by dicarboxylates^3^, requires a low oxygen environment to protect nitrogenase^10^ and involves downregulation of ammonia assimilation into glutamate in bacteroids^18^. Both model predictions and metabolic flux analysis of *A. caulinodans* bacteroids indicate that these factors are interconnected: catabolism of dicarboxylates induces a highly reduced redox poise and creates a high oxygen demand. In the low oxygen environment of the nodule, this promotes downregulation of the decarboxylating TCA cycle arm, which would decrease ammonia assimilation into glutamate by bacteroids. This agrees with glutamate levels being 20-fold lower in bacteroids compared to free-living rhizobia^8^. In addition to promoting ammonia secretion to the plant, limited ammonia assimilation into glutamate would contribute to the growth arrest of bacteroids. As glutamate is the transamination donor for most other amino acids, this could explain the dependence of bacteroids on supply of amino acids by the plant^49,50^. Intriguingly, the increased oxygen stress on bacteroids and their limited ability to assimilate ammonia into glutamate may promote ammonia secretion and could therefore be an important mechanism for maintaining a mutualistic relationship with the host plant. Furthermore, the ample supply of reducing equivalents would then be available to satisfy the high electron demand of the nitrogenase reaction, while carbon polymer synthesis as well as secretion of alanine, in particular, balance the allocation of carbon and regeneration of electron carriers.

The proposed metabolic principles provide a general framework for understanding the constraints on bacteroid metabolism that favour ammonia (and amino acid) secretion and may contribute to the growth arrest of bacteroids. However, rhizobium-legume symbioses are highly evolved mutualisms. As a result, many factors, such as nodule cysteine-rich (NCR) peptides, play a role in inducing terminal differentiation of some bacteroids^51^ and ammonia secretion by bacteroids is further forced by transcriptional regulation of genes involved in nitrogen metabolism (e.g. Ntr)^52^.

Previous modelling studies of inter-microbial interactions found anoxic conditions to promote mutualistic interactions and increase the diversity of secreted metabolites^53,54^, and oxygen availability was determined to be a better indicator of secreted metabolites than species identity^54^. It is therefore possible that metabolic principles similar to those outlined in this study have a broader significance in governing metabolite exchanges in inter-species interactions.

## Methods

### General modelling procedures

Standard FBA calculations were performed in MATLAB R2019b (Mathworks) using scripts from the COBRA Toolbox v3.0^55^ and the Gurobi 8.0.1 solver (www.gurobi.com). The Taxicab norm was minimised to avoid loops in the calculated flux distribution. The functions `phenotypePhasePlane` and `fluxVariability` were used for phenotype phase plane analysis and flux variability analysis, respectively. For flux variability analysis, loopless solutions were calculated that allowed for maximum nitrogenase activity. *In silico* gene deletion analysis was performed with the function `singleGeneDeletion` using the FBA method. A gene was considered essential when its deletion prevented flux through the nitrogenase reaction (Supplementary Data 6). All MATLAB scripts are available on Github (https://github.com/CarolinSchulte/bacteroid-metabolism). Flux maps were created using Escher^56^.

### Elementary conversion mode analysis

Elementary conversion modes were calculated using ecmtool (https://github.com/tjclement/ecmtool)^32^. Boundary conditions for calculating conversion modes were based on experimental studies and are detailed in Supplementary Data 8. To limit our analysis to biologically meaningful scenarios, only conversion modes with a positive cost lower than 40 g carbon per g nitrogen were considered. We further restricted our analysis to conversion modes using one amino acid at most as an input. The full set of conversion modes calculated with succinate, malate and amino acids as inputs can be found in Supplementary Data 7. For determining the correlation of oxygen demand and glutamine synthetase activity, glutamate was set as an additional output and a virtual metabolite was added to the glutamine synthetase reaction to track the flux through this reaction. ECM analysis for *i*CC541, a metabolic model of *Sinorhizobium fredii* bacteroids^19^, was performed according to similar principles. To decrease computation time, cofactors were hidden in this analysis (Supplementary Data 8).

### Statistical analysis

Nonparametric Spearman correlation and ANOVA followed by Tukey’s multiple comparisons test were performed in GraphPad Prism 8.4.3. *P* < 0.05 was considered statistically significant.

### Flux balance analysis

Constraints for FBA-based computations were defined similar to those for conversion mode analysis. For all proteinogenic amino acids except for asparagine, alanine and aspartate, demand fluxes were constrained to values between 0.01 and 0.05 to mimic low levels of protein synthesis in bacteroids. The upper bound for alanine and aspartate demand reactions was left unconstrained, since those amino acids are secreted by bacteroids. All constraints are detailed in Supplementary Data 8.

Ensemble-evolutionary FBA was performed as previously described^47,57^. Briefly, 50,000 objective functions containing a random number of model reactions that were assigned random weights were generated (see Supplementary Fig. 9 for determination of ensemble size). Flux distributions for all objective functions were then calculated using flux balance analysis for a fixed malate uptake rate of 4 flux units and varying oxygen uptake rate. Constraints for ensemble-evolutionary FBA were defined as for standard FBA, but with a minimum flux of 0.01 for the nitrogenase reaction and no upper bound on all amino acid demand reactions.

### Model reconstruction

A database of gene-protein-reaction associations was derived from genome annotation of *Rhizobium leguminosarum* bv. *viciae* 3841 (Rlv3841). First, an orthology-based reconstruction was obtained using AuReMe^58^. Since both a genome-scale model (*i*GD1575b^21,59^) and a highly curated core model (*i*GD726^59^) for the rhizobial strain *S. meliloti* 1021 are available, these were used as templates. Reconstruction in AuReMe was performed for both *S. meliloti* models, resulting in draft networks containing 1304 and 613 reactions for *i*GD1575b and *i*GD726, respectively. Where the same reaction occurred in both reconstructions, the gene-protein-reaction association from the *i*GD726-based reconstruction was selected as it can be expected to be more accurate. To account for gene functions annotated in the Rlv3841 genome but not present in *S. meliloti*, a second draft metabolic model was obtained from KBase^60^. Briefly, the Rlv3841 genome was annotated with Prokka (v1.12) and the ‘Build Metabolic Model’ function was run without gap-filling. This generated a network with 607 reactions, 98 of which were not contained in the reaction list obtained from the AuReMe reconstructions. A model of bacteroid metabolism in Rlv3841 was then built by manually selecting reactions from the draft reconstructions that are catalysed by enzymes detected in the bacteroid proteome, as well as those associated with upregulated^34^ and essential genes for bacteroids^35^. Gene-protein-reaction associations were refined by comparison to gene essentiality data for free-living Rlv3841^61^. Gaps were filled based on literature evidence and guided by the KEGG database^62^. Since the focus of this study was on central metabolic pathways, biosynthesis of some cofactors was excluded and GTP was replaced with ATP where applicable. Some linear pathways, namely most reactions associated with lipid biosynthesis and *myo*-inositol catabolism, were summarised in lumped reactions. Further details on pathways included in the model and the definition of exchange fluxes are provided in Supplementary Note 2 and 3.

### Bacterial strains and culture conditions

Oxygen consumption and NADH/NAD^+^ ratios were measured for Rlv3841. Cultures were grown in universal minimal salts (UMS)^61^ or acid minimal salts (AMS)^63^ medium at 28 °C. UMS was supplemented with different carbon and nitrogen sources at the following final concentrations: succinate, 20 mM; arabinose, 20 mM; glucose, 10 mM; NH_4_Cl, 10 mM. Cultivations were performed in 250 mL Erlenmeyer flasks with an initial filling volume of 50 mL. Low oxygen cultivations were performed in a glove box (Belle Technology) with the atmosphere adjusted to the desired oxygen concentration by flushing with nitrogen gas. *Azorhizobium caulinodans* ORS571 was grown in UMS medium supplemented with 10 mM succinate and 0.3 mM nicotinic acid at 37 ⁰C. For diazotrophic growth, cultures were continuously sparged with a gas mixture containing 97% N_2_ and 3% O_2_, which had been determined to be the optimal oxygen level for diazotrophic growth.

### Measurement of NADH/NAD+ ratios

NADH/NAD^+^ ratios were determined using the NAD/NADH-Glo™ Assay (Promega) according to the manufacturer’s instructions. Cells were harvested during exponential phase (OD_600_ 0.4-0.6) for all measurements.

### Measurement of oxygen consumption rates

Oxygen consumption rates were measured for liquid cultures in early exponential phase. Cultures were diluted to OD_600_ ~0.15 and a 25 mL glass vial containing an OxyDot was filled to the top with liquid culture and sealed. Measurements of oxygen concentration were taken every 15 s using the OxySense 325i system while the culture was stirred with a magnetic stirrer. The data were analysed with the OxySense Gen III software and oxygen consumption was calculated from measurements obtained between 18% and 15% oxygen concentration.

### Proteomics

Bacteroids were obtained from nodules of pea (*Pisum sativum* cv. Avola) plants inoculated with Rlv3841 and grown in an illuminated environment-controlled growth room as previously described^8^. Plants were harvested at 28 d post inoculation, with bacteroids extracted from excised root nodules by double percoll gradient purification^14^. Bacteroids were isolated from nodules obtained from a total of 21 plants and processed as 3 independent replicates.

For the free-living cultures, Rlv3841 was grown in AMS medium with 20 mM succinate and 10 mM ^15^NH_4_Cl. Cultures were grown to late log phase (OD_600_ ~0.8) at 28 °C on a gyratory shaker at 250 rpm, and sub-cultured into fresh AMS six times, as preliminary trials showed this yielded >99% ^15^N incorporation into cell proteins. From the sixth subculture, three separate AMS cultures were inoculated and harvested at mid-log phase (OD_600_ ~0.4). Cells were harvested by centrifugation, washed in AMS and stored at −80 °C. Bacteroid and cell pellets were resuspended separately in 10 mM HEPES (pH 7.2) buffer and an aliquot of each taken and lysed by two rounds on a FastPrep FP120 Ribolyser (BIO101/Savant) at a setting of 6.5 for 30 s, with samples kept on ice for 5 min between each round. The protein content of the extracted aliquots was then determined by Bradford assay, where bovine serum albumin (Pierce™) was used to generate a standard curve. These values were then used to mix equivalent proportions of unextracted bacteroid and cells from the original suspensions to yield at least 200 μg/ml of total combined protein. Mixed bacteroid and cell samples were then ribolysed as described above, another Bradford determination was performed to confirm the protein concentration, and then the equivalent of 50 μg of protein was extracted by methanol-chloroform-water precipitation^64^.

The protein pellets from the three replicates were dissolved in SDS-gel sample loading buffer, heated at 80 °C for 10 min, and loaded onto a Novex-gel (10% Bis-Tris SDS-gel, Life Technologies, Carlsbad, CA). After separation and staining with InstantBlue™ (Expedeon, Harston, UK), the gel lanes were cut into 12-15 slices which were washed, reduced and alkylated, and treated with trypsin according to standard procedures. After digestion, peptides were extracted with 5% formic acid/50% acetonitrile, dried, and re-dissolved in 0.1% TFA. LC-MS/MS analysis of all gel fractions was performed on an LTQ-Orbitrap™ mass spectrometer (Thermo Fisher, Waltham, MA) coupled with a nanoAcquity™ UPLC™-system (Waters, Manchester, UK). Sample aliquots were loaded onto a trap column (Symmetry^R^ C18, 5μm, 180 μm x 20 mm, Waters), and the peptides were then separated on an analytical column (BEH C18, 1.7 μm, 75 μm x 250 mm, Waters) and infused into the mass spectrometer via a 10 μm SilicaTip™ nanospray emitter (New Objective, Woburn, MA) attached to a nanospray interface (Proxeon, Odense, Denmark).

For separation the following gradient of solvents A (0.1% formic acid in water) and B (0.1% formic acid in acetonitrile) was used at a flow rate of 250 nL min^−1^: solvent B at start: 0%; 0-3 min: linear ramp to 5% B; 3-56 min: ramp to 40% B; 56-62 min: ramp to 85% B; 85% B kept for 3 min followed by 100% A for 20 min for re-equilibration.

The mass spectrometer was operated in positive ion mode at a capillary temperature of 200 °C. The source voltage and focusing voltages were tuned for the transmission of MRFA peptide (m/z 524) (Sigma-Aldrich, St. Louis, MO). Data-dependent analysis was carried out in orbitrap-IT parallel mode using CID fragmentation on the five most abundant ions in each cycle. The full scan MS was performed in the orbitrap at a resolution of 60,000 over the range *m/z* 350-1800. For the CID-MS2 the mono-isotopic 2+ and 3+ charged precursors were selected with an isolation width of 2 Da. MS2 was triggered by a minimal signal of 10^3^ with an AGC target of 3×10^4^ ions and 150 ms maximum scan time using the chromatography function for peak apex detection. Collision energy was 35, and dynamic exclusion was set to 1 count and 60 s with a mass window of ±20 ppm. MS scans were saved in profile mode while MS2 scans were saved in centroid mode.

The analysis of all gel fractions from 3 replicates resulted in 52 raw files (with replicate 2 run twice). Data were processed using Mascot Distiller 2.7 and Mascot Server 2.7 through Mascot Daemon 2.7 (Matrixscience, London, UK). Peak lists generated by Mascot Distiller were used for a database search using Mascot Server on the GeneDB_Rleguminosarum_Proteins fasta database from https://www.sanger.ac.uk/resources/downloads/bacteria/rhizobium-leguminosarum.html (May 2020, 7144 entries). For protein annotation, data from https://rhizosphere.org/lab-page/molecular-tools/genomes/rlv3841-genome/ (May 2020, 7288 entries) was used. A small database containing common contaminants (MaxQuant contaminants 2017, 250 entries) was included in the search.

Each raw file was processed and searched separately using the enzyme trypsin with two missed cleavages, precursor mass tolerance of 10 ppm and fragment mass tolerance of 0.6 Da. Carbamidomethyl (C) was set as a fixed modification, and oxidation (M), deamidation (N,Q), acetylation (protNterm) as variable modifications. Intensities for light and heavy ^15^N labelled peptides were extracted using a Mascot Server ^15^N metabolic quantitation method with the following parameters: 99.2% labelling (as determined by Mascot Distiller), Simpson’s integration, isotope match rho = 0.6, XIC threshold = 0.1, isolation threshold = 0.5, peptide expect threshold = 0.05, outlier removal auto, normalization none. The experimental design with 3 replicates and corresponding fractions was generated in the quantitation table exported via Mascot Daemon. The resulting expression table was used for ratio calculation and statistical analysis in RStudio 1.2.5033 with R version 3.6.3^65^. Potential contaminants were removed, and the table was filtered for proteins quantified in all 3 replicates. Light and heavy protein intensities were log10-transformed, and ratios calculated as light/heavy (bacteroids/*Rhizobium* cells) for each replicate. Those ratios were used for statistical testing using the limma eBayes function in R. The final ratio was calculated as median from the replicates.

The mass spectrometry proteomics data have been deposited to the ProteomeXchange Consortium via the PRIDE^66^ partner repository with the dataset identifier PXD019467.

### Metabolic flux analysis

Metabolic flux analysis of *A. caulinodans* ORS571 was performed as previously described^8^. Briefly, *A. caulinodans* was grown in UMS medium containing 20% [^13^C_4_]-succinate and 80% unlabelled sodium succinate. Bacteroids were isolated from the root nodules of *Sesbania rostrata* as previously described^67^. Isolated bacteroids were incubated in UMS medium supplemented with 10 mM 20% [^13^C_4_]-succinate for 24 h at ≤1% O_2_. Nitrogenase activity of free-living cultures and isolated bacteroids during labelling experiments was determined by an acetylene reduction assay^8^. Free-living cultures were harvested in late exponential phase and bacteroids after 24 h for metabolite extraction and gas chromatography-mass spectrometry analysis. Mass spectrometry data processing and isotopomer analysis of protein-derived amino acids and hydroxybutyrate obtained by hydrolysis of PHB was done using methods described previously^8,68^.

Metabolic modelling was performed with INCA (Isotopomer Network Compartmental Analysis) using an iterative procedure^69^. A complete description of the model, including the network carbon atom transitions, and net flux data are provided in Supplementary Data 9. The model along with the mass spectrometry measurements was simulated to obtain an optimized flux pattern in the network, followed by statistical validation of the flux maps. The fluxes were estimated relative to the succinate uptake flux fixed at 100 units. The assessment of goodness of fit for the flux maps with statistically valid sum of squared residuals was performed by the comparison of simulated mass spectrometry measurements with that of experimentally measured values. To assess the precision of flux estimates, parameter continuation was performed in INCA to calculate the lower and upper bounds of the 95% confidence interval for the flux estimates. Comparison of fluxes estimated for the free-living bacteria and bacteroids were based on the confidence intervals (upper and lower limits) for a specific flux. A flux determined under two different conditions was deemed to be significantly different if the confidence intervals for the flux under the two conditions did not overlap.

## Data availability

All data needed to evaluate the conclusions in this paper are present in the main text or supplementary materials. Proteome data are available via ProteomeXchange with identifier PXD019467. All code is available on Github (https://github.com/CarolinSchulte/bacteroid-metabolism).

## Acknowledgements

The authors would like to thank Prof. Bas Teusink at VU Amsterdam for helpful discussions and Carlo de Oliveira Martins at the John Innes Centre for support with analysing the proteome data. Chiara Damiani at University of Milano-Bicocca is acknowledged for providing the ensemble-evolutionary FBA scripts.

This work was supported by funding from the Biotechnology and Biological Sciences Research Council [grant numbers BB/F013159/1, BB/M011224/1] and Natural Environment Research Council [grant number NE/L501530/1]. D.H.d.G. was supported by NWO VICI grant 865.14.005. A.P. was funded in part by the Engineering and Physical Sciences Research Council [grant number EP/M002454/1]. C.C.M.S is supported by the Clarendon Fund (Oxford University Press) and the Keble College De Breyne Scholarship. K.B. was supported by the Louis Dreyfus Weidenfeld Scholarship for Plant Science from the University of Oxford.

## Author Contributions

C.C.M.S. performed model reconstruction and analysis, redox ratio and oxygen consumption measurements. K.B. performed metabolic flux analysis. R.M.W. performed preliminary experiments and contributed to conception of the study. J.J.T. and G.S. performed proteome quantification. N.C. performed experiments supporting the interpretation of the metabolic flux analysis data. D.H.d.G. assisted with the design and implementation of the conversion mode analysis. All authors analysed the data. P.S.P, A.P., N.J.K. and R.G.R. designed and supervised the study. C.C.M.S. wrote the manuscript, with input from K.B., J.J.T., G.S., N.J.K., R.G.R., A.P. and P.S.P.

## Materials and correspondence

The datasets generated and analysed during the current study are available from P.S.P. and N.J.K. upon request.

