## Supplementary Information for "Metabolic constraints on nitrogen fixation by rhizobia in legume nodules"

**Supplementary Note 1. Evaluation of bacteroid proteome and transcriptome data.** The proteome of bacteroids was compared with that of free-living *Rhizobium leguminosarum* bv. *viciae* 3841 (Rlv3841) growing on minimal media supplemented with succinate and $^{15}NH_4Cl$ as the sole carbon and nitrogen source, respectively. A total of 1087 proteins were identified and grouped according to their Riley classification. 106 proteins had significantly increased abundance in bacteroids, 426 proteins had decreased abundance and 555 were unchanged.

The adaptation to symbiotic conditions was reflected by 49 and 40 proteins involved in adaptation and regulation, respectively (Supplementary Fig. 1a). These were predominantly heat shock proteins and transcriptional regulators. This group also contained enzymes related to glutathione (GSH) metabolism and detoxification of reactive oxygen species (ROS). ROS are produced during the rhizobium-legume symbioses both as signalling molecules and as part of the plant defence response. The ability to detoxify ROS, either enzymatically or through production of antioxidant molecules, such as GSH, is essential for an effective symbiosis[1]. As expected, the components of nitrogenase (Nif proteins) and associated electron transfer proteins (Fix proteins) were highly increased in abundance[2].

Despite the non-growing state of bacteroids, 26.3% of the proteins detected were associated with basic cell functions, such as cell envelope, nucleic acid and protein synthesis and degradation, which is consistent with previous reports of protein secretion by bacteroids[3] and the known phenomenon of DNA endoreduplication[4].

The largest group of proteins detected (34.2%) was broadly classified as metabolic enzymes. Regarding central carbon metabolism, TCA cycle enzymes were identified in accordance with the essentiality of this pathway in Rlv3841 bacteroids[5]. In addition, enzymes of the pentose phosphate and Entner-Doudoroff pathway were found[6,7]. The presence of several respiratory chain enzymes reflects the obligate aerobic metabolism even under low oxygen conditions inside a nodule[8]. 79 proteins were involved in amino acid biosynthesis, with 22 of these being downregulated compared to free-living bacteria. This result indicates active synthesis of proteins as well as amino acid exchange with the plant host as demonstrated previously[9,10]. Further enzymes important for amino acid metabolism were found in the group of intermediary metabolism. Notably, this group contained gabT and gabT2, two enzymes of the $\gamma$-aminobutyric acid (GABA) pathway, which is highly induced in pea bacteroids[11].

The role of storage polymers in rhizobium-legume symbioses has been reported[5,12], and proteins associated with polyhydroxybutyrate (PHB) (phaA, phaB, phaC, phaE), glycogen (glgC) and lipid (various Fab proteins) synthesis were detected. Several proteins involved in *myo*-inositol catabolism (IolABCDE) were present in agreement with the high concentrations of *myo*-inositol in nodules[13], even though mutants in inositol catabolism did not show a symbiotic phenotype in Rlv3841[14].

Finally, 8.6% of the detected proteins were putatively involved in transport processes. Apart from the broad-specificity amino acid transporters Aap and Bra[15] with known importance for the symbiosis[9,16], most of the other transport proteins are poorly characterized in Rlv3841.

Among the 106 proteins showing significantly increased abundance in bacteroids, the Nif and Fix proteins were highly upregulated, along with cytochrome oxidases (Supplementary Fig. 1b). Proteins involved in PHB synthesis were highly abundant, emphasising the importance of this storage polymer, which had initially been thought to be absent from pea bacteroids[5].

Apart from GABA metabolism mentioned above, a homoserine O-acetyltransferase was strongly upregulated among the enzymes involved in amino acid metabolism. Starting from homoserine, an abundant and probably plant-provided metabolite in bacteroids[5], this enzyme catalyses the first step towards methionine synthesis. HisI, a protein involved in histidine biosynthesis, also showed highly increased abundance. Together with the presence of five other His proteins (HisB, HisC1, HisD1, HisH, HisF), this indicates active biosynthesis of histidine in bacteroids, which agrees with studies on histidine auxotrophs of *Rhizobium meliloti* that could not be rescued by the plant host[17]. A propionate CoA-transferase (pRL100119) also showed significantly increased abundance. This gene has high similarity to general acyl CoA:acetate/3-ketoacid CoA-transferases in various bacteria and is thus likely involved in lipid metabolism.

To further support the selection of pathways to include in the metabolic model, data from a previously published transcriptome study of Rlv3841 bacteroids[2] were evaluated. We limited our analysis to the 136 genes found to be at least 3-fold upregulated in pea bacteroids at 28 d post inoculation relative to the free-living state. A large fraction of upregulated genes (34.6%) could not be assigned a function (Supplementary Fig. 2a). The second major difference to the proteome data was the decreased fraction of genes associated with general cell functions (14% compared to 26.3% in proteome data). This difference is due to the consideration of all detected proteins, not just upregulated proteins, in the proteome data analysis. Cell growth- and maintenance-associated proteins are expected to be decreased in expression due to the non-growing state of the bacteroid, and their abundance was decreased accordingly in the proteome data.

The genes/proteins identified in both datasets supported the inclusion of PHB, methionine and histidine metabolism. Further upregulated genes identified in the transcriptome dataset (Supplementary Fig. 2b) included a phenylalanine-4-hydroxylase, consistent with significant labelling of phenylalanine in bacteroids of pea plants incubated in $^{15}N_2$[11]. An acetyl-CoA synthetase (pRL100121) showed a 4-fold increase in expression. This gene has high similarity to a characterized *acsA1* gene in *S. meliloti*[18] and is an important connection between anabolic and catabolic reaction.

**Supplementary Note 2. Pathways included in *i*CS323.**

*Central carbon metabolism*: The TCA cycle was included as the main route for metabolising dicarboxylic acids provided by the plant. Activity of all TCA cycle enzymes has been detected in bacteroids of *R. leguminosarum*[19], the genes have been found to be transcriptionally upregulated and corresponding proteins identified in the proteome data. Enzymes involved in glycolysis and gluconeogenesis, such as pyruvate kinase (PykA) and phosphoenolpyruvate carboxykinase (PckA), were detected in the bacteroid proteome in accordance with previous studies[6] and insertions in the *pckA* as well as fructokinase (*frk*) genes had deleterious effects on the symbiosis[20]. Even though these pathways are expected to be less important in bacteroids than dicarboxylic acid metabolism, they are required e.g., for nucleotide and glycogen synthesis. *myo*-inositol is abundant in pea nodules[13] and may be an additional carbon source provided by the plant. All proteins encoded by the *iolABCDE* genes were detected in bacteroids.

*Amino acid metabolism*: Despite the non-growing state of bacteroids, protein turnover is still occurring, as evidenced by labelling studies[11] and the detection of ribosomal subunits in the proteome. It is important to account for metabolism of amino acids provided by the plant host[9,16] and synthesis of the exported amino acids alanine and aspartate[10,21]. According to the experimental datasets used in this study, homoserine O-acetyltransferase was one of the most highly upregulated enzymes in bacteroids. Homoserine is probably provided by the plant host[5] and functions as a precursor for methionine synthesis. Multiple proteins involved in histidine, arginine and tryptophan biosynthesis were detected in bacteroids. Biosynthetic pathways for all amino acids except for asparagine, which is not synthesized in its free form by Rlv3841[5], were therefore included in the model. In addition, GABA catabolism, which has been shown to be strongly induced in bacteroids compared to free-living bacteria[11], was included.

*Carbon polymers*: Pathways for the biosynthesis of PHB and glycogen were included since both polymers are produced by bacteroids of *R. leguminosarum*[5,12] and especially proteins involved in PHB synthesis showed significantly increased abundance in bacteroids. Lipids have further been identified as an important carbon sink in bacteroids[5], and several Fab proteins involved in lipid metabolism were identified in the proteome. A representative species for free fatty acids and glycerolipids was therefore included in the model.

*Other polymers*: Homospermidine and polyphosphate have been shown to be present in bacteroids[22,23] and pathways for their synthesis were added to the model.

**Supplementary Note 3. Nutrient exchange between pea plants and *R. leguminosarum* bacteroids.** Based on a review of experimental evidence, we defined the boundary conditions for nutrient exchange between bacteroid and plant host.

*L-Malate:* It is well established that dicarboxylic acids, such as malate, succinate and fumarate, are the main carbon sources fuelling symbiotic nitrogen fixation. A recent study found that malate is sufficient as the main carbon source to support efficient nitrogen fixation by Rlv3841[24].

*L-Leucine:* Transport of branched-chain amino acids is required at low levels to support an efficient symbiosis, with leucine being the most likely to be required from the plant host[16].

*Homoserine:* Homoserine levels are high in pea bacteroids[5] and although catabolic genes (*hom1*, *hom2*) were not upregulated, a putative homoserine O-acetyltransferase (MetX, encoded by pRL100137) was highly increased in abundance compared to free-living cells, indicating an important role for homoserine as a precursor for amino acid synthesis.

*Other amino acids*: While the requirement for branched-chain amino acids has been explicitly demonstrated[16], this does not preclude provision of other amino acids by the plant. For example, enzymes involved in GABA metabolism are highly active in bacteroids, even though this pathway does not seem to be essential for efficient nitrogen fixation[11]. Elementary conversion mode analysis was therefore performed for all proteinogenic amino acids and GABA as inputs.

While it is generally accepted that ammonia is the main nitrogenous compound supplied to the plant by the bacteroid, experimental evidence suggests that amino acids such as alanine and aspartate may play a role as additional secretion products[10,25]. Demand reactions for those amino acids were therefore also included.

**Supplementary Table 1. Properties of the experimental datasets and the bacteroid model *i*CS323.**

| **Bacteroid datasets** | |
|---|---|
| Total identified proteins | 1087 |
| Involved in metabolism | 489 |
| Included in model | 218 |
| Upregulated genes | 136 |
| Involved in metabolism | 49 |
| Corresponding protein detected | 52 |
| Included in model | 28 |
| Essential genes for bacteroids and nodule bacteria | 258 |
| Involved in metabolism | 69 |
| Corresponding protein detected | 70 |
| Gene upregulated | 11 |
| Included in model | 33 |
| ***i*CS323** | |
| Genes | 323 |
| Metabolites | 237 |
| Unique EC identifiers | 176 |
| Reactions | 299 |
| Metabolic reactions | 207 |
| Gene-associated metabolic reactions | 203 |
| Transport reactions | 32 |
| Exchange and demand reactions | 60 |
| Metabolic reactions supported by experimental evidence | 177 |

a

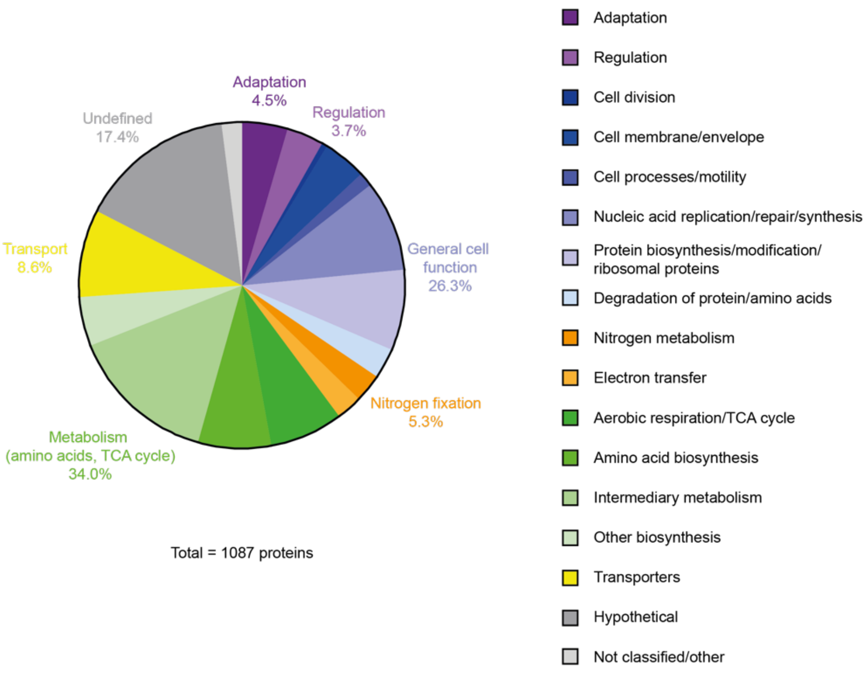

b

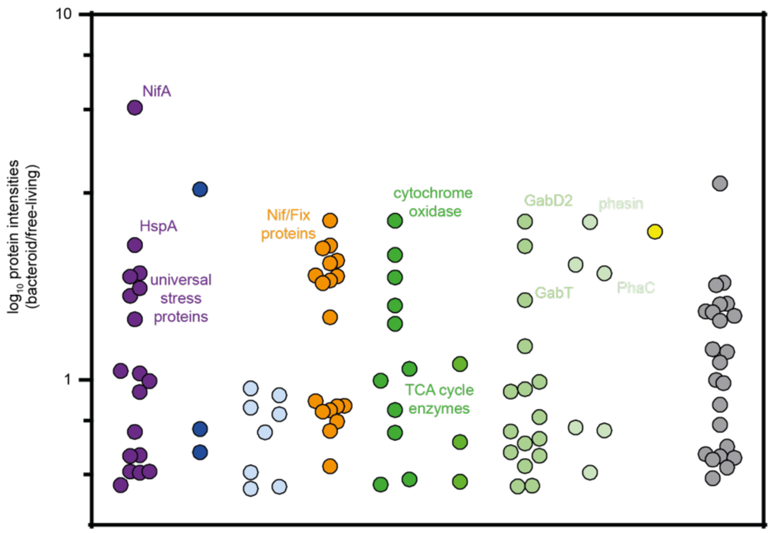

**Supplementary Figure 1. Comparative proteomics of bacteroids and free-living Rlv3841.** The proteome of bacteroids at 28 d post inoculation and free-living bacteria grown on succinate and ammonia was compared. **a** Pie chart showing the functional distribution of all proteins detected in bacteroids according to their Riley classification. **b** Scatter plot showing the median of the log10 ratio bacteroid/free-living for the intensities of the 106 proteins that were significantly increased in abundance in bacteroids. Proteins are grouped along the x-axis according to their Riley classification.

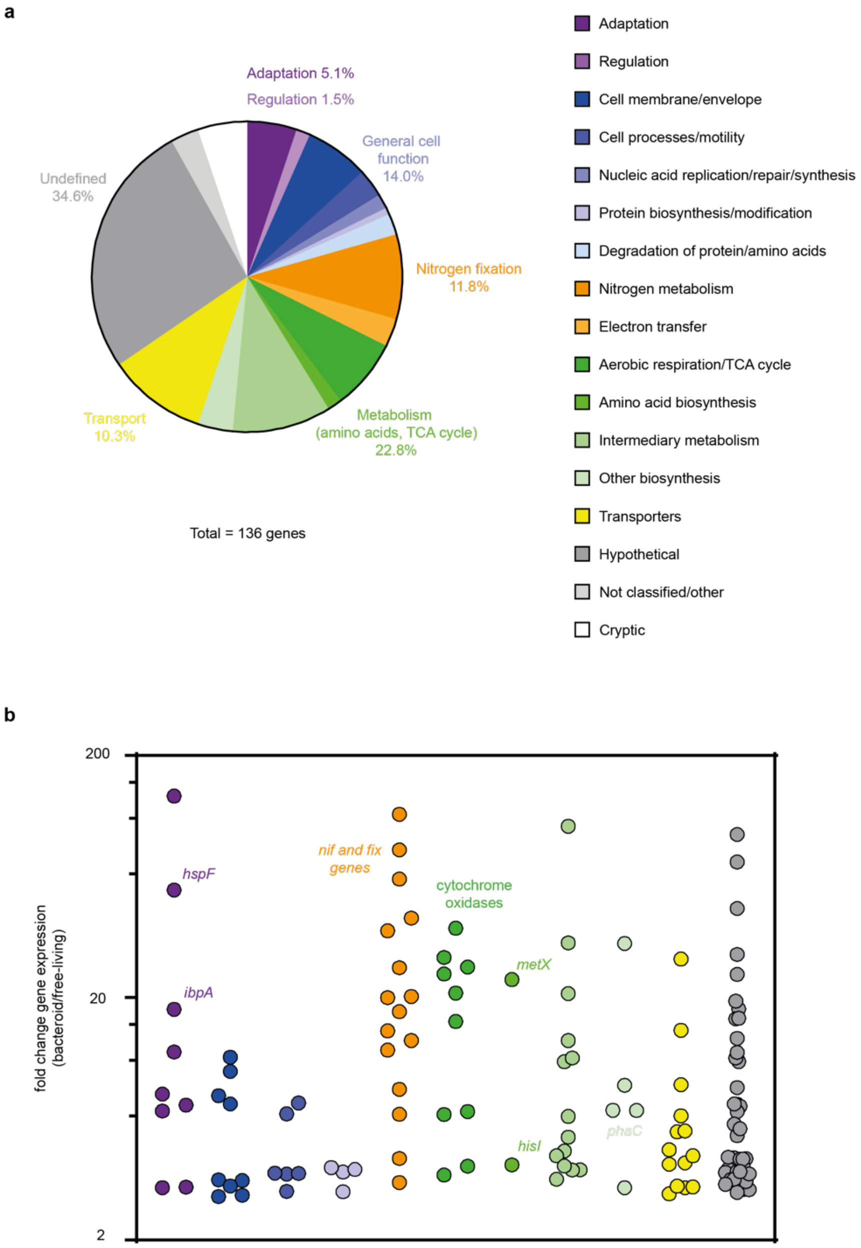

**Supplementary Figure 2. Comparative transcriptomics of bacteroids and free-living Rlv3841**. The transcriptome of bacteroids at 28 d post inoculation and free-living bacteria grown on succinate and ammonia[2] was compared. Only genes with at least 3-fold upregulation in bacteroids are shown. **a** Pie chart showing the functional distribution of genes upregulated in bacteroids according to their Riley classification. **b** Scatter plot showing the fold-change (bacteroid/free-living) in expression for genes upregulated in bacteroids. Genes are grouped along the x-axis according to their Riley classification.

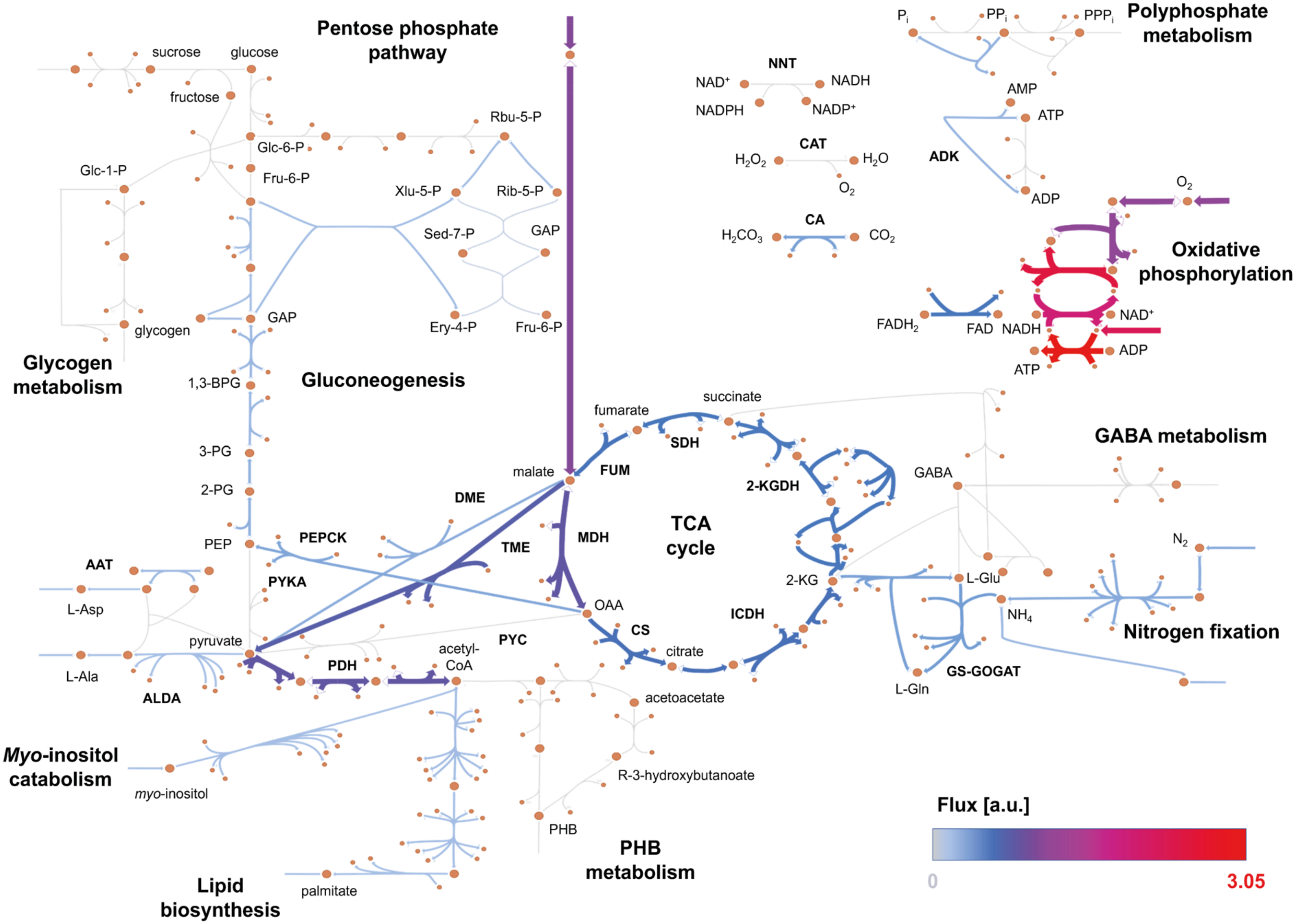

Pentose phosphate pathway
Polyphosphate metabolism
Glycogen metabolism
Gluconeogenesis
Oxidative phosphorylation
TCA cycle
GABA metabolism
Nitrogen fixation
Myo-inositol catabolism
Lipid biosynthesis
PHB metabolism
sucrose
glucose
fructose
Glc-6-P
Fru-6-P
Glc-1-P
glycogen
Rbu-5-P
Xlu-5-P
Rib-5-P
Sed-7-P
GAP
Ery-4-P
Fru-6-P
1,3-BPG
3-PG
2-PG
PEP
pyruvate
L-Asp
L-Ala
AAT
ALDA
PEPCK
PYKA
DME
TME
MDH
FUM
SDH
2-KGDH
ICDH
CS
PYC
PDH
malate
fumarate
succinate
OAA
citrate
2-KG
acetyl-CoA
acetoacetate
R-3-hydroxybutanoate
PHB
palmitate
myo-inositol
GABA
L-Glu
L-Gln
$NH_4$
$N_2$
GS-GOGAT
NNT
$NAD^+$
NADH
NADPH
$NADP^+$
CAT
$H_2O_2$
$H_2O$
$O_2$
CA
$H_2CO_3$
$CO_2$
$P_i$
$PP_i$
$PPP_i$
AMP
ATP
ADK
ADP
$FADH_2$
FAD
Flux [a.u.]
0
3.05

**Supplementary Figure 3. Summary map of *i*CS323, a model of bacteroid metabolism in Rlv3841.** A metabolic network of pea bacteroids was reconstructed from proteome, transcriptome and gene essentiality data. The map summarises reactions in the model with colours corresponding to a flux distribution for maximum nitrogenase activity with malate, succinate and oxygen uptake rates constrained according to boundary conditions used in a modelling study of *Sinorhizobium meliloti* bacteroids[26]. Colour and arrow thickness correspond to flux values with thin grey arrows indicating inactive reactions. Some pathways for amino acid biosynthesis have been omitted for clarity.

AAT: aspartate aminotransferase, ADK: adenylate kinase, ALDA: alanine dehydrogenase, CA: carbonic anhydrase, CAT: catalase, CS: citrate synthase, DME: NAD-dependent malic enzyme, FUM: fumarase, GS-GOGAT: glutamine synthetase-glutamine oxoglutarate aminotransferase, ICDH: isocitrate dehydrogenase, 2-KGDH: 2-ketoglutarate dehydrogenase, MDH: malate dehydrogenase, NNT: nicotinamide nucleotide transhydrogenase, PYC: pyruvate carboxylase, PDH: pyruvate dehydrogenase, PEPCK: phosphoenolpyruvate carboxykinase, PYKA: pyruvate kinase, SDH: succinate dehydrogenase, TME: NADP-dependent malic enzyme; 1,3-BPG: 1,3-bisphosphoglycerate, Ery-4-P: erythrose 4-phosphate, Fru-6-P: fructose 6-phosphate, GAP: glyceraldehyde 3-phosphate, Glc-1-P: glucose 1-phosphate, Glc-6-P: glucose 6-phosphate, 2-KG: 2-ketoglutarate, OAA: oxaloacetate, 2-PG: 2-phosphoglycerate, 3-PG: 3-phosphoglycerate, PEP: phosphoenolpyruvate, PHB: polyhydroxybutyrate, Rib-5-P: ribose 5-phosphate, Rbu-5-P: ribulose 5-phosphate, Sed-7-P: sedoheptulose 7-phosphate, Xlu-5-P: xylulose 5-phosphate

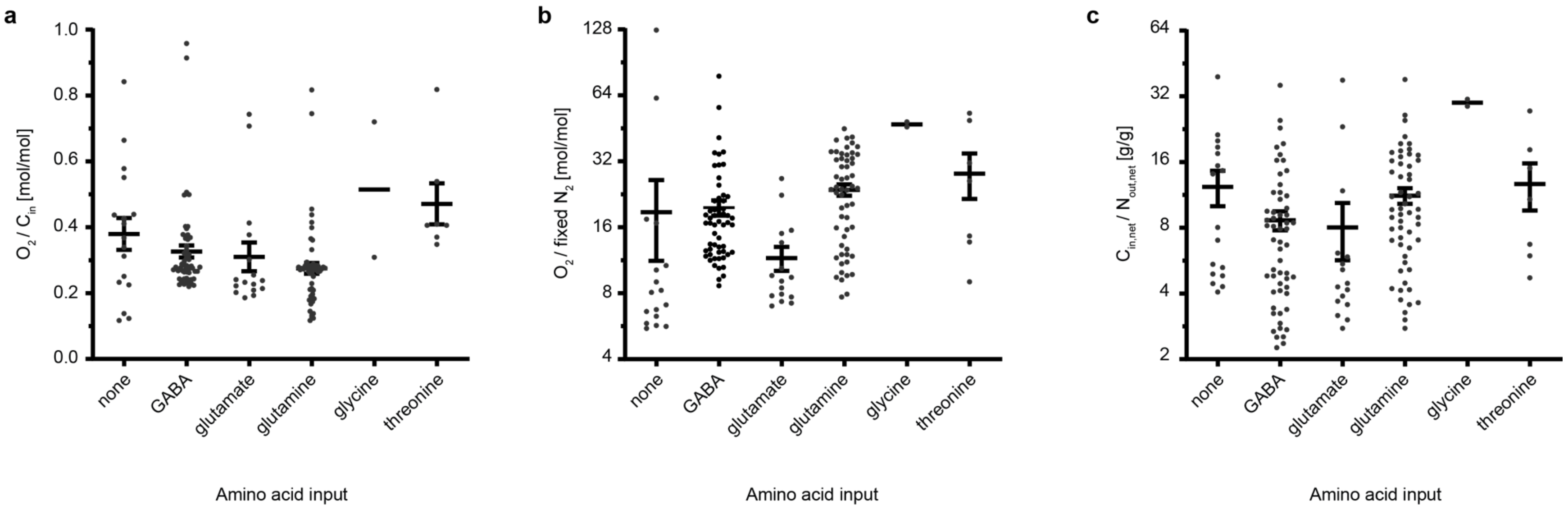

**Supplementary Figure 4. Effect of different amino acid inputs on oxygen demand and carbon cost of nitrogen fixation.** Elementary conversion modes were calculated with malate, succinate and different amino acids as inputs and carbon polymers (palmitate, PHB, glycerolipid, glycogen), ammonia, alanine and aspartate as outputs. **a** Oxygen uptake per carbon uptake, **b** oxygen uptake per fixed $N_2$ and **c** carbon cost (difference of carbon input and output) per nitrogen secreted (difference of nitrogen input and output). Conversion modes are grouped according to the amino acid in the input. Each data point represents an individual conversion mode, lines and bars indicate mean ± SEM.

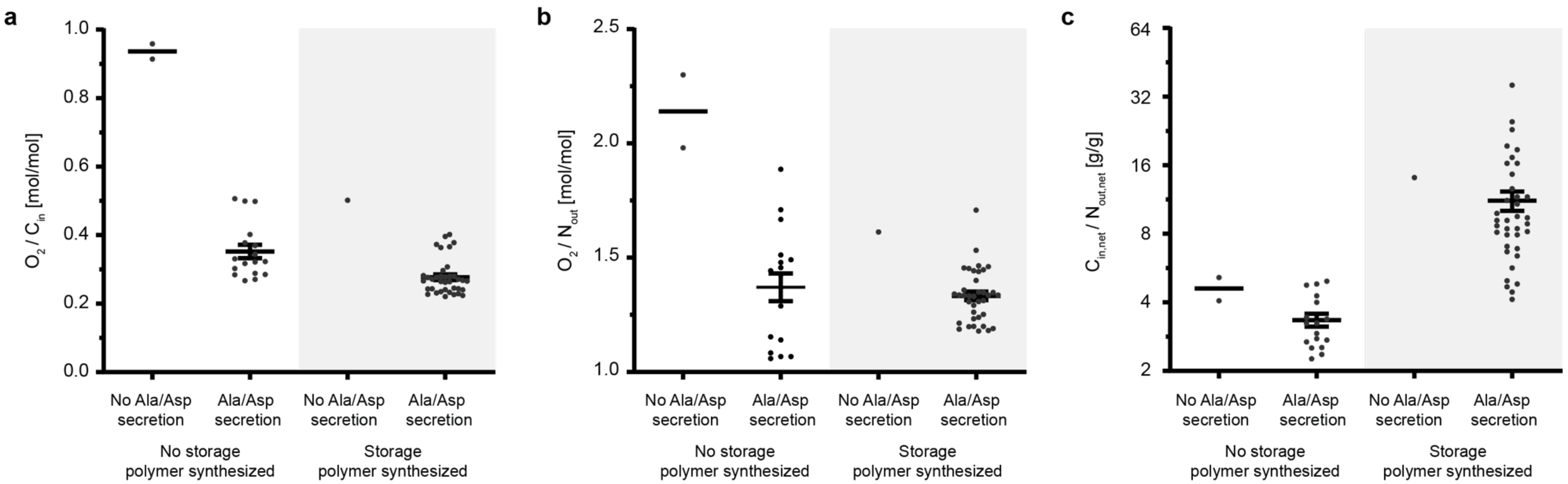

**Supplementary Figure 5. Effect of polymer synthesis and amino acid secretion on oxygen demand and carbon cost of nitrogen fixation.** Elementary conversion modes were calculated with malate, succinate and GABA as carbon sources and carbon polymers (palmitate, PHB, glycerolipid, glycogen), ammonia, alanine and aspartate as outputs. **a** Oxygen uptake per carbon uptake, **b** oxygen uptake per nitrogen output and **c** carbon cost (difference of carbon input and output) per nitrogen secreted (difference of nitrogen input and output). Each data point represents an individual conversion mode, lines and bars indicate mean ± SEM.

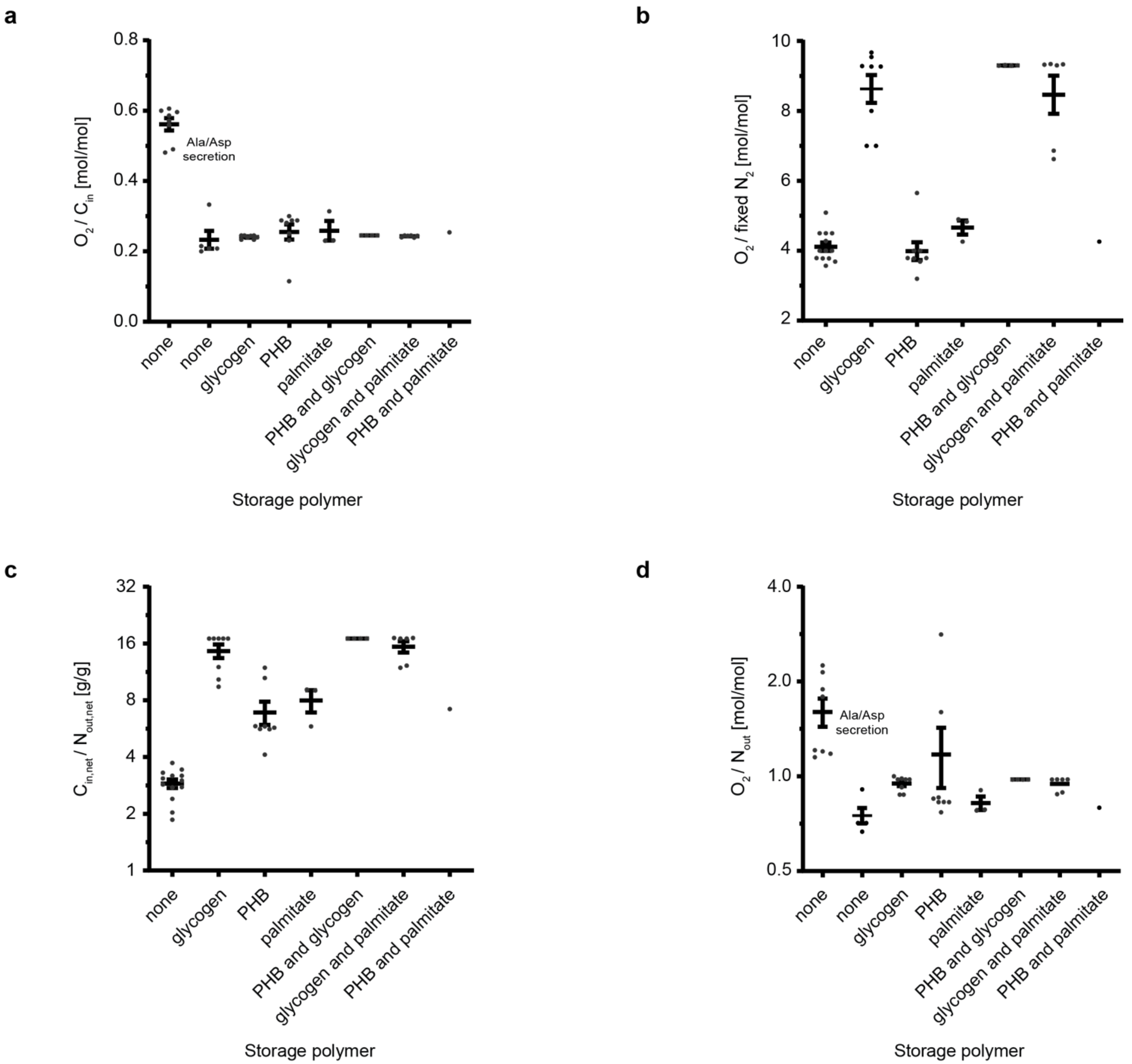
a
O2 / Cin [mol/mol]
0.0
0.2
0.4
0.6
0.8
Ala/Asp secretion
none
none
glycogen
PHB
palmitate
PHB and glycogen
glycogen and palmitate
PHB and palmitate
Storage polymer
b
O2 / fixed N2 [mol/mol]
2
4
6
8
10
none
glycogen
PHB
palmitate
PHB and glycogen
glycogen and palmitate
PHB and palmitate
Storage polymer
c
Cin,net / Nout,net [g/g]
1
2
4
8
16
32
none
glycogen
PHB
palmitate
PHB and glycogen
glycogen and palmitate
PHB and palmitate
Storage polymer
d
O2 / Nout [mol/mol]
0.5
1.0
2.0
4.0
Ala/Asp secretion
none
none
glycogen
PHB
palmitate
PHB and glycogen
glycogen and palmitate
PHB and palmitate
Storage polymer

**Supplementary Figure 6. Elementary conversion mode analysis of *Sinorhizobium fredii* bacteroids.** Conversion modes for the metabolic model *i*CC541[27] were calculated with succinate, malate, and glutamate as the main carbon inputs and carbon polymers (PHB, glycogen, palmitate), ammonia, alanine and aspartate as outputs. **a** Oxygen uptake per carbon uptake, **b** oxygen uptake per fixed $N_2$, **c** carbon cost (difference of carbon input and output) per nitrogen secreted (difference of nitrogen input and output) and **d** oxygen uptake per nitrogen output. In **a** and **d**, ECMs without storage polymer production have been separated into those secreting only ammonia and those secreting alanine and/or aspartate in addition to/instead of ammonia. Each data point represents an individual conversion mode, lines and bars indicate mean ± SEM.

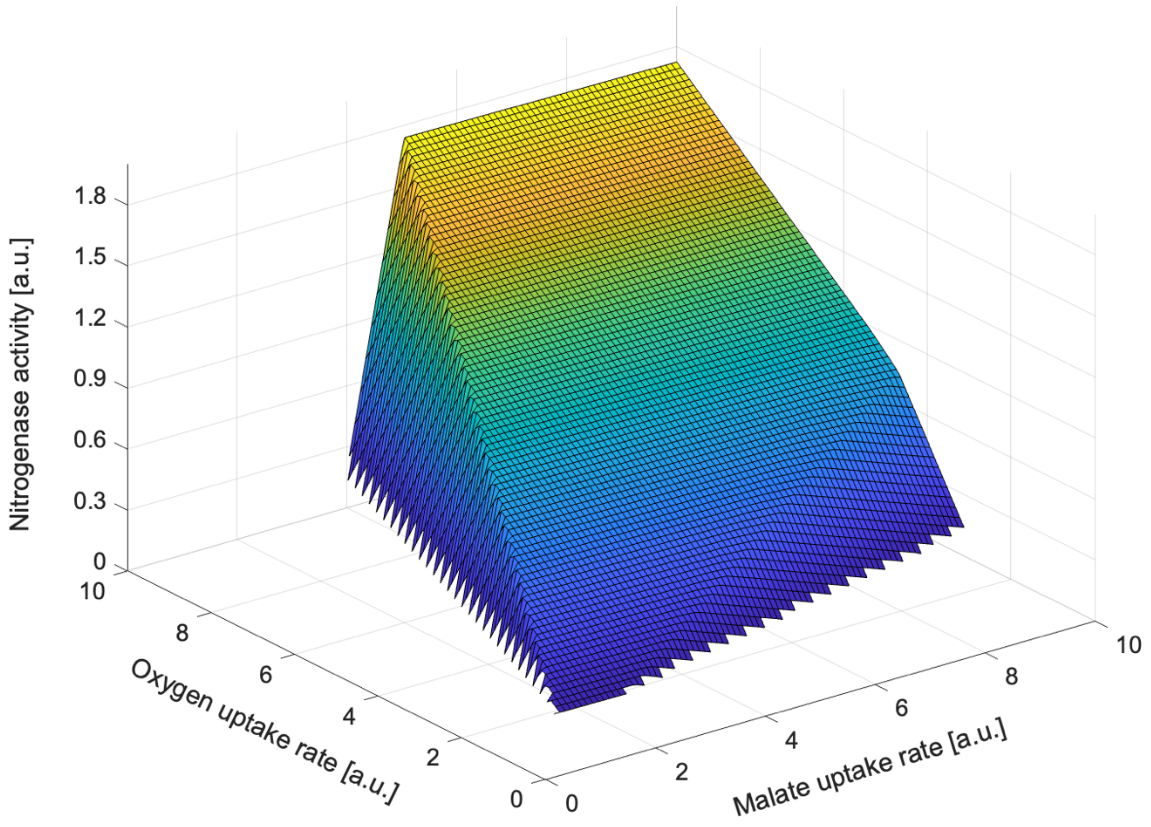

**Supplementary Figure 7. Phenotype phase plane analysis of *i*CS323.** Three-dimensional representation of the phenotype phase plane analysis of *i*CS323. Colours correspond to nitrogenase activity.

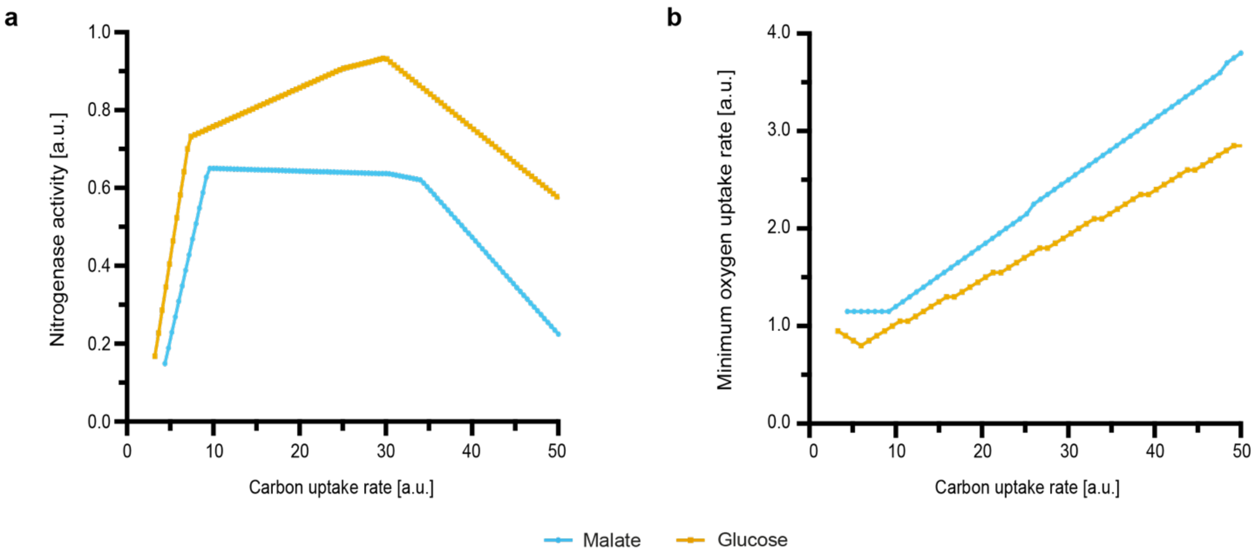

**Supplementary Figure 8. Predicted effect of malate and glucose catabolism on nitrogenase activity and oxygen demand. a** Maximum nitrogenase activity for malate or glucose as a carbon source with maximum oxygen uptake of 4 flux units. **b** Minimum oxygen demand for nitrogenase activity with malate or glucose as a carbon source. Values are shown per mol of carbon uptake.

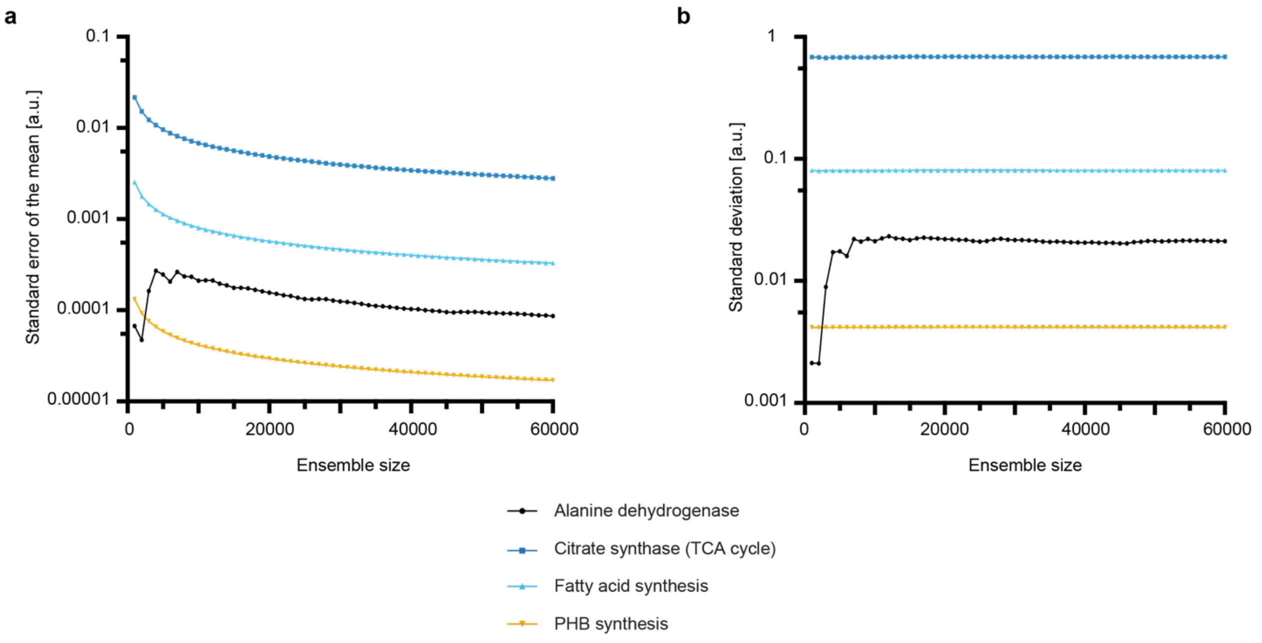

**Supplementary Figure 9. Determination of ensemble size for ensemble-evolutionary flux balance analysis.** To define the ensemble size (number of random objective functions) used in ensemble-evolutionary flux balance analysis computations, **a** the standard error of the mean and **b** the standard deviation for different ensemble sizes were calculated for the reactions of interest.
